# Analysis of 3.6 million individuals yields minimal evidence of pairwise genetic interactions for height

**DOI:** 10.1101/2024.08.15.608197

**Authors:** M.Reza Jabalameli, Michael V Holmes, David Hinds, 23andMe Research Team, Adam Auton, Pierre Fontanillas

## Abstract

Adult height is a highly heritable polygenic trait with heritability attributable to thousands of independent variants. Large-scale studies have been able to detect genetic variants with contributions to height in the range of approximately 1.2 millimetre per allele copy on average. Non-additive genetic interactions may, in part, account for the difference between broad-sense and narrow-sense heritability estimates. However, prior studies have failed to identify variants with non-additive effects, possibly due to the lack of statistical power. Leveraging 3.6M individuals of European genetic ancestry in the 23andMe research cohort, we performed a genome-wide analysis study (GWAS) to select 1,063 independent common SNPs associated with height (p-value < 5e-8), and then screened for evidence of non-additive effects by analysing 564,453 models including a pairwise SNP-SNP interaction term. We identified 69 pairwise models with suggestive evidence of SNP-SNP interaction (p-value < 1e-4) and, for each SNP pair, we evaluated a fully saturated model including additive, dominant, and epistatic (additive-by-additive, additive-by-dominance and dominance-by-dominance) terms. We tested for the presence of epistatic interactions by comparing models with and without epistatic terms using a likelihood ratio test. Assuming a strict Bonferroni-corrected threshold of 8.9e-8 (0.05/564,453), we found no evidence of epistatic interactions (Likelihood ratio test (LRT) p-value < 9e-07 for all models). Our analysis rules out the existence of epistatic interactions between alleles of >1% frequency with effect sizes larger than 2.42mm. Our large-scale analysis provides further evidence of the minimal contribution of non-additivity in the genetic architecture of adult human height.

## Introduction

Adult height is largely determined by genetics, with about 80% of its variation inherited through genetics^1^. However, the role of non-additive gene actions, such as pairwise genetic interaction (G×G), in determining height is still unclear, with past studies showing minimal impact^2^, potentially due to limited statistical power.

Genetic interactions are historically studied in inbred model organisms where the phenotype of double mutants is compared against single mutants in an identical genetic background to identify genes acting in the same pathway or convergent molecular functions^3^. While there are numerous examples of genetic interactions in model organisms^4^, evidence for genetic interactions in human traits is sparse^5^.

Height is reliably measured, enabling large sample sizes which should provide sufficient statistical power to detect genetic interactions. GWAS analyses of height have revealed that the trait is highly polygenic, involving thousands of common variants with mean effect size estimates in the range of approximately 1.2 millimetre per allele copy^6,7^. The most recent GWAS of height^7^ identified 7,209 loci spanning 647Mb of the genome (∼ 21%) to explain approximately >94% of common (MAF > 1%) SNP-based heritability in Europeans. While the identified common variant associations saturate common SNP-based heritability, they only explain 40% of the interindividual variation in height among Europeans. This estimate is less than the total heritability of 80% reported among identical and fraternal twins^8^. The missing heritability in height has been attributed to a number of factors^9^, including imperfect linkage disequilibrium (LD) between the tagged SNPs and the actual causal variants, rare variants that are not accurately imputed, and overestimation of the heritability due to gene-by-gene and gene-by-environment interactions^10^. Analysis of variants under the GWAS framework relies on the “additive mean effect”, where individual variants are assumed to affect the mean trait value independently of other variants. Conversely, genetic variants associated with trait variance (i.e interindividual variation that is under genetic control^11^) have been reported for quantitative traits such as body mass index (BMI)^12^ and blood glucose level^13^.

Epistasis is a biological phenomenon wherein the influence of one genetic variant on a complex trait is contingent upon the genotype of a second genetic variant (which manifests as deviation from additivity with respect to model of additive effects^14^) and is suspected to play an important role in the genesis of complex disease^15^. Deviation from additivity can either arise from “dominance” or “genetic interactions” between variants. The distinction between the two is that in dominance, the interaction is between alleles within the same locus, while in epistasis, interaction occurs between different loci. Simulation analysis of pairwise interactions has shown that the power to replicate the main effect of a variant dramatically diminishes as allele frequency at the second interacting loci drops below 10%^16^. Non-additive gene action (i.e. the specific way a gene affects an organism’s phenotype that is not additive) is well documented in model organisms, but in humans, despite the dramatic increase in sample size of biobanks, evidence for non-additivity is sparse due to the lack of sufficient power for detecting epistatic interactions.

Population biobanks of millions of individuals offer the opportunity to explore the contribution of epistasis to human complex traits. Here, using data from 3.6 million individuals from the 23andMe, Inc. research cohort, we set out to investigate the evidence of pairwise genetic interactions in height. Testing for all pairwise interactions is computationally intensive and incurs a considerable penalty for multiple testing corrections; therefore, we implemented a two-stage strategy^17–19^ to investigate pairwise interactions among GWAS variants with significant additive effects. In testing pairwise interactions, we fully partition the non-additive component to identify true additive-by-additive interactions that are independent of dominance and dominance-by-additive interactions (**Figure 1**).

**Figure 1:**
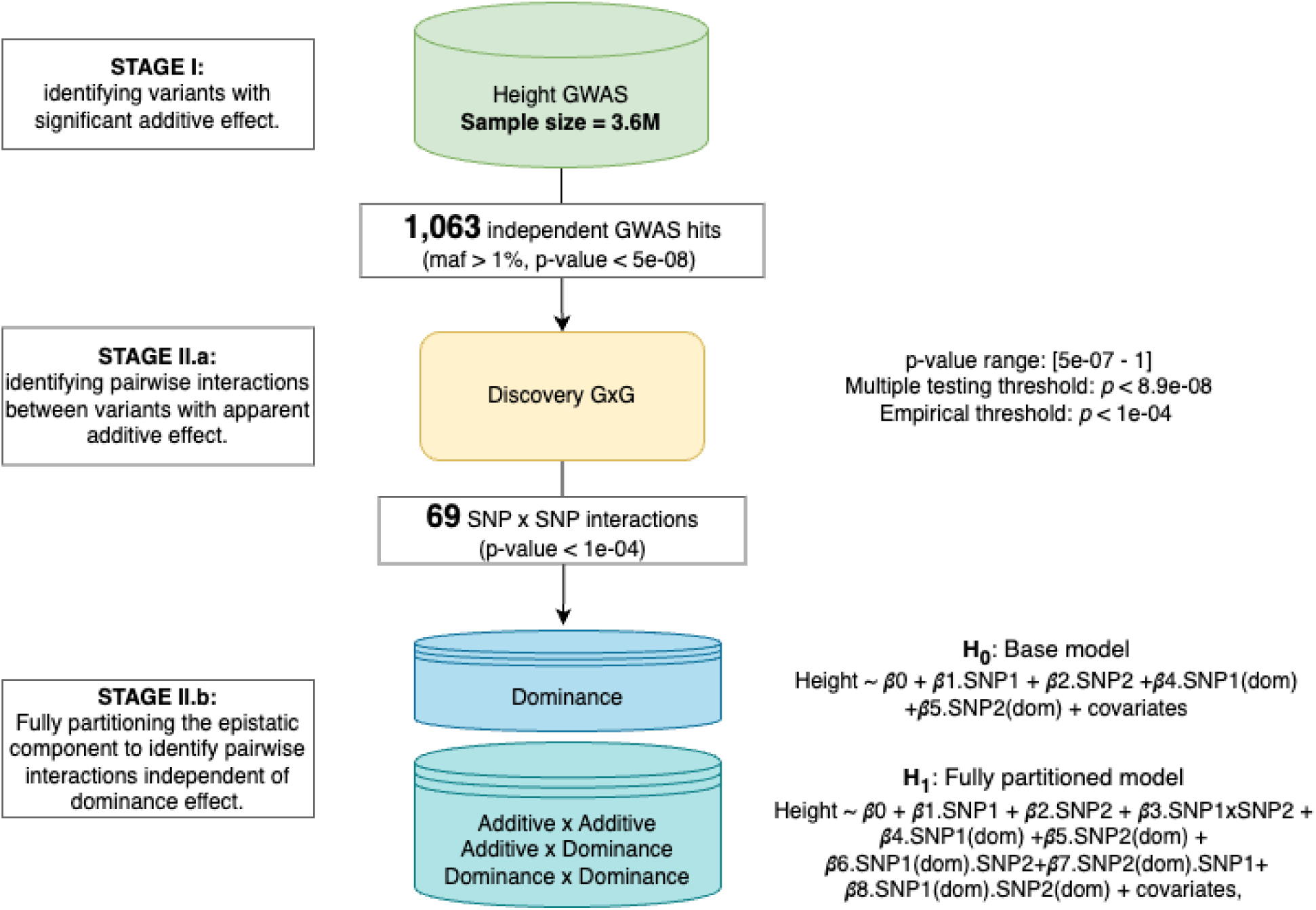
Study design and analysis schema. We followed a two-stage approach to identify interactions between pairs of genetic variants. First, we conducted a GWAS of height using data from approximately 3.6 million participants (Stage I). From this analysis, we selected 1,063 independent variants that met the genome-wide significance threshold (p < 5e-08) to investigate potential pairwise interactions. We tested 564,453 pairwise interactions and found 69 pairs that exceeded our empirical threshold (p < 1e-04) (Stage II.a). We then employed two linear models (LMs) to test for epistatic interactions between these 69 SNP pairs and their potential dominance effect.

## Results

### GWAS of adult height and pairwise GWAS SNP-SNP interaction screen

A total of 3,652,184 participants were included in the GWAS analysis of adult height. Distribution of individuals across quantiles of height measurements, stratified by sex and age groups is provided (male: µ = 179.80 cm, sd= 7.03; female: µ= 164.83 cm, sd= 6.77) (**Supplementary Table 1**). The association of 15,698,169 variants that passed the QC process was investigated using linear regression while adjusting for sex, age, top five principal components and genotyping platforms (**Materials & Methods**). We applied LD score regression (LDSC) to quantify the genomic inflation and calculate heritability (h^2^=0.38, intercept=3.77). We found 1,063 independent index SNPs with MAF >1% surpassing the genome-wide significance threshold (p < 5e-8), with an average estimated absolute effect of 1.71 millimetres (**Supplementary Table 2** and **Supplementary Figure 1 & 2**). The minimum observed distance between two independent associations was approximately 509.22kb (**Supplementary Figure 3 and Materials & Methods**).

For the 1,063 identified index variants, we screened for evidence of non-additive effects by adding a pairwise SNP-SNP interaction term to the original GWAS model. In total, we tested *564,453* pairwise interactions, of which none surpassed the multiple testing corrections 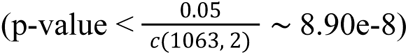. We further removed any interactions within 2Mb of each other (495 tests). The distribution of interaction p-values and QQ plot is provided in **Supplementary Figure 4**. Recognising the stringency of the Bonferroni correction, which may be overly conservative in the context of genetic interactions, we adopted a more lenient p-value threshold of < 1e-04 for identifying putative interactions. This approach aligns with previous studies^20^ that have utilised the same (or more lenient^21^) thresholds to balance the detection of true signals against the risk of false positives. We identified 69 SNP pairs (involving 138 unique index variants) with suggestive evidence of (G×G) interaction (p-value < 1e-4) (**Supplementary Table 3** and **Supplementary Figure 5**). These putative interactions include six intrachromosomal interactions (median distance 26.8 Mb) and 66 interchromosomal hits. With only one exception, the absolute magnitude of the interaction effect size between *intrachromosomal*-interactions was smaller than the additive term effects from either interacting loci (one-sided signed binomial test (***H_0_*** : β_*GxG*_ > β_*additive*_) p-value= 0.11, 95% CI=0.0 - 0.58; **Supplementary Figure 6**). Similarly, the general trend among *interchromosomal*-interactions followed the same pattern as *intrachromosomal*-interactions, where the absolute magnitude of effect sizes across the 66 interactions were smaller than the individual additive term effect size of either interacting loci (one-sided signed binomial test p-value = 2.2e-16, 95% CI=0.0 - 0.06, **Supplementary Table 4**).

We observed that in both *intrachromosomal* and *interchromosomal*-interactions, the direction of interactions was mainly discordant with the additive effects of either one or both interacting loci. Out of the total interactions, we found only one concordant *intrachromosomal*-interaction and 11 concordant *interchromosomal*-interactions (**Supplementary Figure 6** and **Supplementary Table 3)**. We did not identify any specific pattern (such as proximity to regulatory elements or TF binding sites) that could determine the consistency of the direction of the interaction effect with the additive effect.

### Fully saturated model with additive and non-additive components

Recognizing that dominance effects within gene pairs might obscure or falsely suggest SNP interactions, we adopted a nested model approach. We fitted two complementary models testing the pairwise interactions while fully partitioning the dominance and epistatic components (**Figure 1, Materials & Methods**). We exclusively modelled dominance across the 69 SNP pairs (with suggestive evidence of interaction) to investigate whether detected interaction is the result of dominance spillover from either interacting loci^22^. Of the 138 SNPs included in our pairwise tests, we identified five variants with significant dominance effects at our lenient threshold (p < 1e-4). Among these variants, only one variant (rs11205303) shows significant dominance effect at Bonferroni-corrected threshold (p= 2.6e-16) **(Supplementary Table 5** and **Supplementary Note**). Of the 69 pairs, five pairs had at least some evidence of dominance.

Next, we tested whether pairwise interactions were explained by dominance (**Materials & Methods**). For the 69 SNP-pairs models, we constructed a fully saturated gene action model including additive, dominant, epistatic (additive-by-additive, additive-by-dominant, and dominant-by-dominant), as well as GxE (variant-by-sex, variant-by-age and variant-by-ancestry effects (i.e., the product of individual genotypes and first two genomic PCs) and ExE (sex-by-age) terms. This methodological framework allows us to discern whether SNP interactions exert a significant influence on height beyond the additive and dominance effects of the interacting alleles (**Figure 1, Materials & Methods**). From the fully saturated model, we identified eight interchromosomal interactions at the lenient threshold of p < 1e-4 (**Table 1**). We considered the genes identified in the putative interacting loci, and found no biological plausibility for bona fide interactions. For the 69 pairwise models, the average absolute effects of the additive-by-additive interactions (mean= 0.86 mm; SD= 1.76) were approximately 2.48 times smaller than the average additive effects (mean= 2.13 mm; SD= 3.32) in the saturated model.

**Table 1:** Summary of SNP-pairs exhibiting suggestive additive-by-additive interaction from fully saturated model at Bonferroni-corrected cutoff of *p_GxG_* < 1e-04.

| Interacting SNP-pairs |  |  |  |  |  |  |  |  |  | Interaction statistics |  | LRT |
| --- | --- | --- | --- | --- | --- | --- | --- | --- | --- | --- | --- | --- |
| SNP1 |  |  |  |  | SNP2 |  |  |  |  |  |  |  |
| Nearby Gene | Position (GRCh38) | rsID | EA/NEA | EA freq (%) | Nearby Gene | Position (GRCh38) | rsID | EA/NEA | EA freq (%) | β (SE) | p-value | p-value |
| GRM1 | chr6:145998281 | rs2748497 | T/C | 43.38 | DPP4 | chr2:162074215 | rs13015258 | G/T | 39.12 | 0.39 mm | 4.24E-06 | 1.50E-04 |
|  |  |  |  |  |  |  |  |  |  | -9.00E-02 |  |  |
| MON2 | chr12:62543609 | rs12370246 | T/C | 64.93 | IRS2 | chr13:110026904 | rs9559710 | C/T | 54.74 | -0.42 mm | 1.33E-05 | 1.67E-06 |
|  |  |  |  |  |  |  |  |  |  | -9.00E-02 |  |  |
| SLC44A2 | chr19:10637445 | rs3843751 | T/C | 67.47 | PLEC | chr8:143948489 | rs7832643 | G/T | 40.02 | 0.43 mm | 1.68E-05 | 2.30E-04 |
|  |  |  |  |  |  |  |  |  |  | -9.00E-02 |  |  |
| RPLP1 | chr15:69742660 | rs71393467 | G/A | 9.47 | TUT7 | chr9:86471377 | rs750187 | A/T | 50.9 | -1.25 mm | 1.95E-05 | 3.05E-05 |
|  |  |  |  |  |  |  |  |  |  | -2.90E-01 |  |  |
| NEDD4 | chr15:55836332 | rs10526731 | I/D | 38.1 | ADGRL2 | chr1:82481993 | rs6661091 | A/C | 66.66 | -0.44 mm | 3.16E-05 | 1.30E-04 |
|  |  |  |  |  |  |  |  |  |  | -1.00E-01 |  |  |
| SMIM23 | chr5:171857638 | rs11424823 | I/D | 62.83 | SLC8A1 | chr2:40628415 | rs7606482 | A/G | 57.46 | -0.34 mm | 2.86E-04 | 3.60E-04 |
|  |  |  |  |  |  |  |  |  |  | -1.00E-01 |  |  |
| N4BP2L2 | chr13:32552699 | rs11318593 | I/D | 63.84 | CREB3L2 | chr7:137911401 | rs273961 | C/T | 60.7 | -0.34 mm | 3.05E-04 | 4.30E-04 |
|  |  |  |  |  |  |  |  |  |  | -1.00E-01 |  |  |
| BCL2L2-PA<br>BPN1 | chr14:23310668 | rs3210043 | C/A | 84.51 | SLC25A13 | chr7:96415301 | rs5885961 | D/I | 44.52 | 0.66 mm | 3.61E-04 | 2.50E-04 |
|  |  |  |  |  |  |  |  |  |  | -1.80E-01 |  |  |

The eight putative pairwise interactions are identified in the fully partitioned model (p-value < 1e-04). Each row represents a unique pair of interactions, and the result of the likelihood ratio test (LRT) comparing the goodness of fit between the base model and the fully saturated model is provided in the last column. Genomic coordinates are defined according to GRCh38; * The effect sizes and standard errors (SE) are expressed in millimetres (note that none of these variants surpassed the Bonferroni-corrected significance threshold of p < 8.9e-08); EA: effect allele; NEA: non-effect allele.

### Variance explained by different variables in the saturated model

We evaluated the relative contributions of genetic and non-genetic components to the variance in height in the fully saturated model. The analysis revealed that the majority of the phenotypic variance in height is attributable to additive gene action (**Figure 2**, and **Supplementary Figure 7**). On average the additive genetic effects of a pair of interacting SNPs explains 0.035% (interquartile (IQR) range: 0.010-0.047%, SE=4.4e-3) of the total phenotypic variance, and 41.4 % of the total variance explained by the model (IQR: 23.2-59.1%, SE=2.5). In contrast, the non-additive components, including dominance and various forms of epistatic interactions (additive by additive, additive by dominance, and dominance by dominance), contributed minimally to the phenotypic variance. On average, dominance deviation across the interacting variant pairs explains 1.36e-04% (IQR = 2.02e-05-9.04e-05%, SE=3.40e-5) of the height variance and epistatic interactions explains 5.26e-04% (IQR= 4.75e-04-6.14e-04%, SE= 1.95e-05).

**Figure 2:**
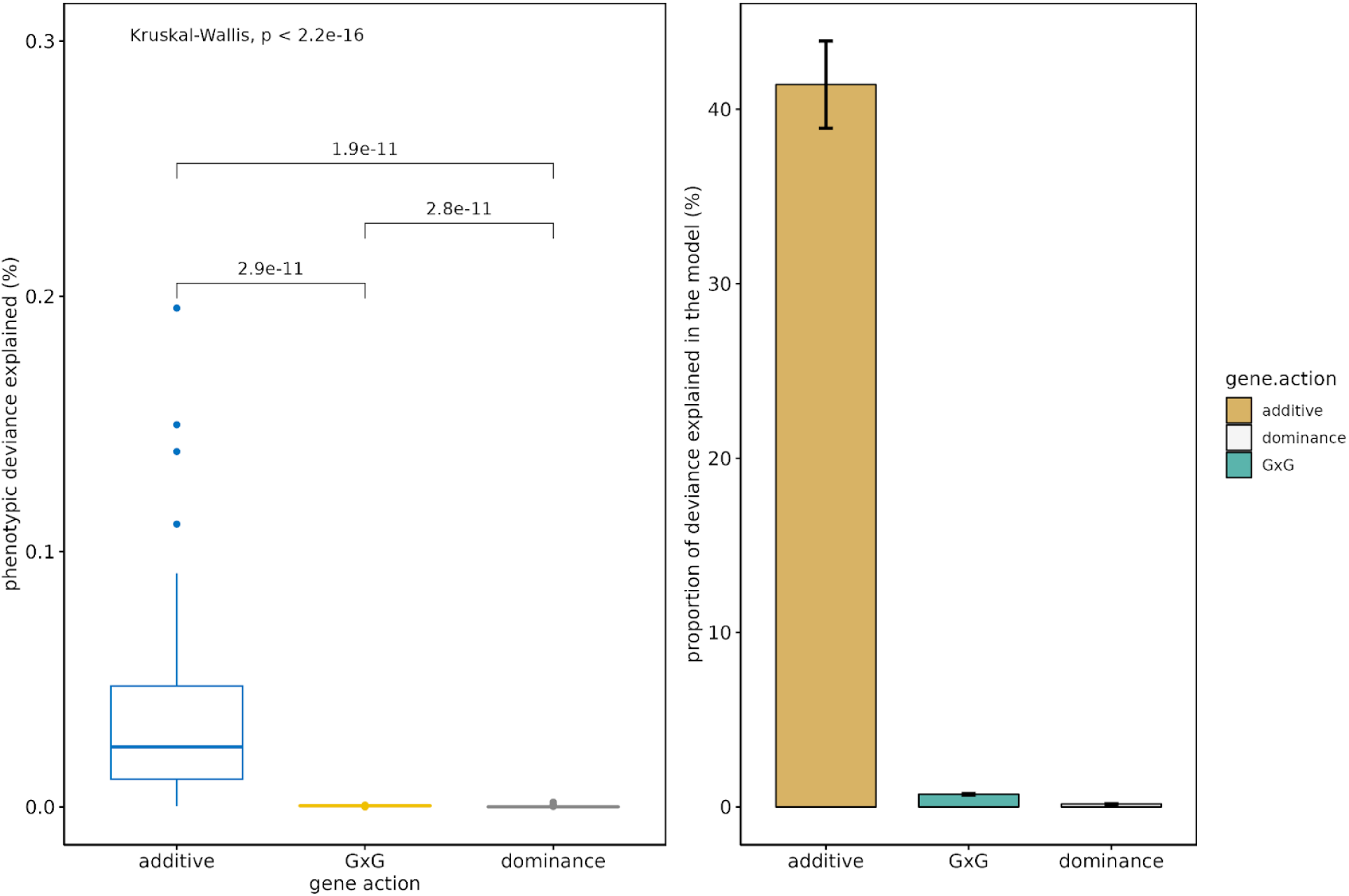
Comparison of various genetic effects’ contributions to height variance in the fully saturated model.

Notably, the variance explained by epistatic interactions is approximately 1/67th or 0.015 folds smaller than additive variance, implying negligible epistatic variance for height. This infinitesimal contribution suggests that while these interactions exist, their effect is such that their impact on height variation is negligible. This finding aligns with the classical understanding of quantitative genetics where additive effects often capture the largest share of variance in polygenic traits^23^. Distribution of deviance explained by additive, dominance and additive by additive gene actions across the 69 tested interaction pairs is provided in **Supplementary Figures 8.a to 8.c)**.

**Figure 2.a (left panel) - Total phenotypic variance explained by various gene actions:** The box plot displays the distribution of the percentage of total phenotypic variance explained by three types of gene actions: additive, additive-by-additive interaction (G×G), and dominance. Each box represents the interquartile range (IQR) of the data, with the central line indicating the median. Outliers are depicted as individual points. A global Kruskal-Wallis test confirms significant differences between the types of gene actions (p < 2.2e-16), with subsequent pairwise comparisons showing significant differences between each category (p-values indicated above the brackets). **Figure 2.b (right) - Proportion of variance explained by gene actions in the model**: The bar plot illustrates the percentage of variance explained by the model that is attributable to each gene action. The height of the bars reflects the mean proportion of variance explained, and the error bars represent the standard error of the mean. Consistent with the box plot, the additive gene action is shown to account for the majority of the variance explained by the model, followed by a marginal contribution from G×G interactions, and the least from dominance.

Comparison of variance explained by different forms of epistatic interactions revealed higher contribution of additive-by-additive, followed by additive-by-dominance and dominance-by-dominance (Figure 3). On average, additive-by-additive genetic effects explain 4.31e-04% (IQR range: 4.23e-04-4.96e-04%, SE= 1.78e-05) of phenotypic variance and only 0.7% of the model’s total variance (IQR: 0.48-0.98%, SE= 0.04). In comparison, additive-by-dominance gene actions accounted for 3.32e-05% (IQR range: 3.69e-06-4.01e-05%, SE= 6.18e-06) of the total phenotypic variance and 0.1% of the total variance explained by the model (IQR range: 0.03-0.15, SE= 0.01). Dominance-by-dominance gene actions had the lowest contributions to the total phenotypic variance (2.87e-05%, IQR range: 1.45e-06-2.73e-05%, SE= 5.66e-06), revealing negligible effect of these epistatic interactions to the total phenotypic variance. These results reaffirm the predominance of additive effects over non-additive interactions in explaining phenotypic variation. The minor role of epistasis, as quantified in our analysis, suggests that its contribution to the genetic variance of height is limited.

**Figure 3:**
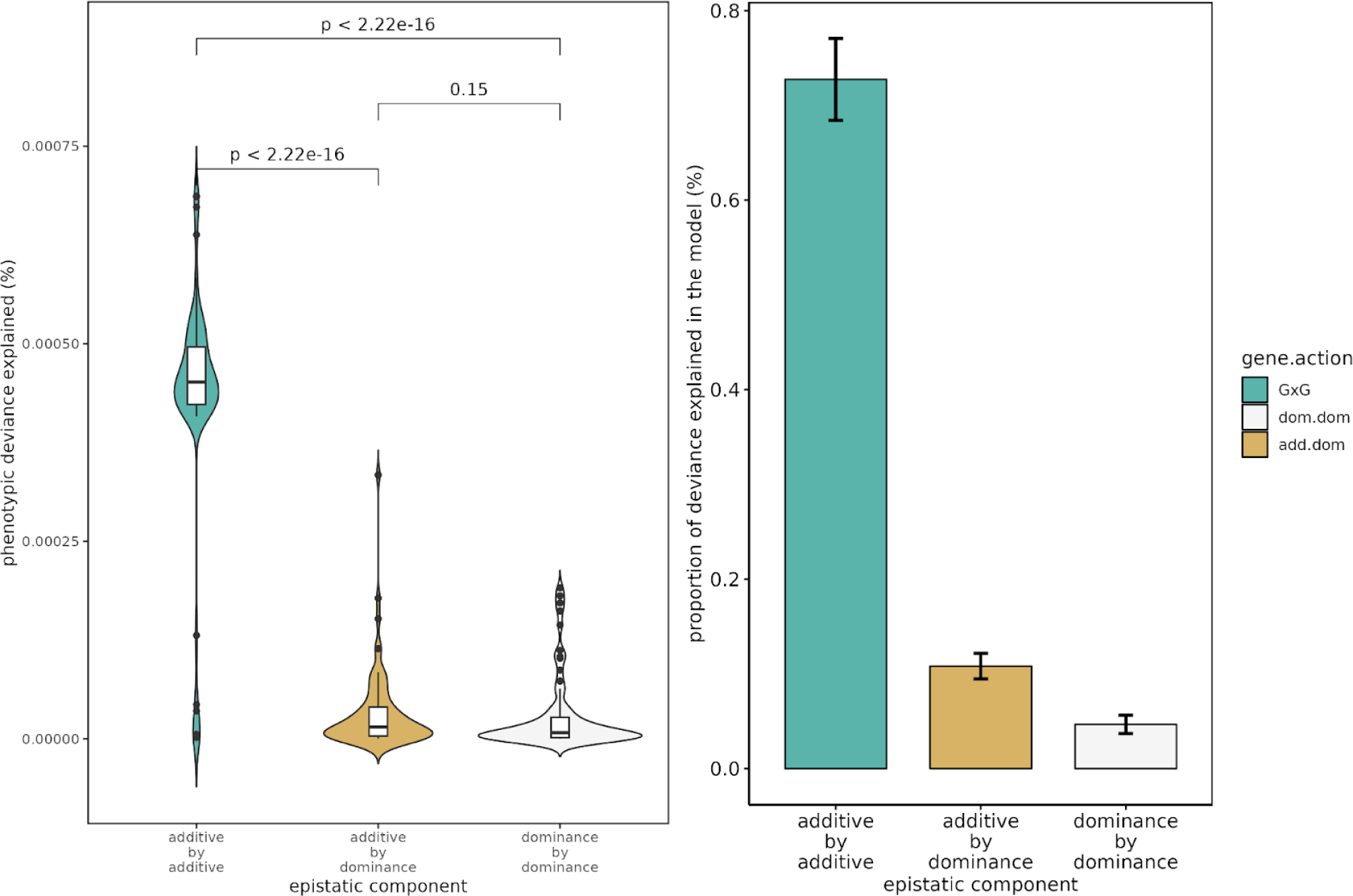
Contribution of various epistatic components to height variance, as explained in the fully saturated model.

We also calculated the variance explained by non-genetic factors and their interactions in the fully saturated model (**Supplementary Table 3**). A detailed explanation of the variance attributed to non-genetic factors is included in the **Supplementary Note**.

**Figure 3.a (left panel) - Total phenotypic variance explained by different epistatic gene actions:**The violin plots paired with internal box plots that depict the distribution of the percentage of total phenotypic variance explained by three types of epistatic interactions: additive-by-additive, additive-by-dominance, and dominance-by-dominance. The shape of the violin plots illustrates the density distribution of the variance explained, with the width corresponding to the frequency of data points. The internal box plots represent the interquartile range (IQR), median, and outliers for each type of interaction. The statistical significance between the types of interactions is denoted by the ‘p-values’ displayed above the comparisons, indicating that there are significant differences in the amount of variance explained by each epistatic component. **Figure 3.b (right panel) - Proportion of explained variance by epistatic interactions in the model:** The bar plot shows the mean proportion of the variance explained by the model that is attributable to each type of epistatic interaction, with error bars representing the standard error of the mean. The height of the bars reveals that additive-by-additive interactions account for a larger proportion of the explained variance, followed by additive-by-dominance, and finally dominance-by-dominance interactions.

Finally, using the theoretical expectation of additive-by-additive effect size range in the GWAS SNP-SNP interaction model, we computed the power to detect a significant pairwise interaction in our 3.6 million cohort. We estimate that we have over 80% power to detect a 1.10mm interaction for allele frequencies greater than 5%, and 80% power to detect a 2.42mm interaction for allele frequencies greater than 1% (Figure 4; **Supplementary Figure 11)**. This suggests that the study’s large sample size is sufficient to ensure a high probability of detecting G×G interactions, even for variants with relatively low allele frequencies.

**Figure 4:**
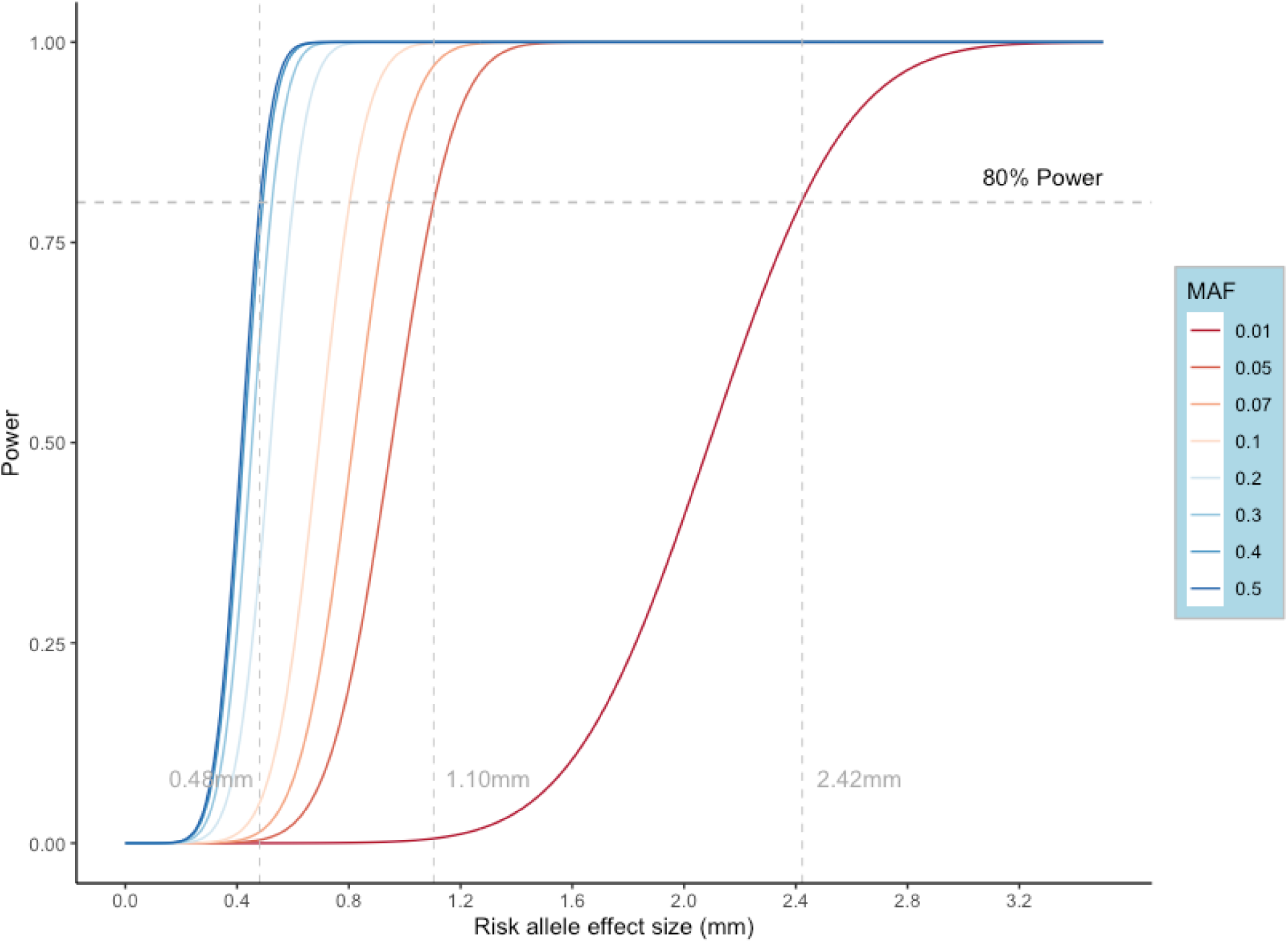
Statistical power to detect (G×G) interaction in the discovery stage (STAGE II.a) as a function of absolute additive-by additive interaction effect size.

This figure illustrates the power to detect additive-by-additive (G×G) genetic interactions across a range of effect sizes and minor allele frequencies (MAFs). Each curve represents a different MAF, from common variants with an MAF of 0.5 (dark blue) to rare variants with an MAF of 0.01 (dark red). The x-axis reflects a continuum of effect sizes in millimetres, and the y-axis denotes the power to detect interactions. The dashed horizontal lines represent critical power thresholds, with the upper line marking the conventional 80% power level considered sufficient for reliable detection. The curves reveal that for variants with an MAF of 0.5, the study possesses high power across virtually all observed effect sizes. As the MAF decreases, the curves shift to the right, indicating that a larger effect size is necessary to achieve the same level of power.

## Discussion

We looked for evidence of pairwise interactions in adult height across 3.6 million participants of the 23andMe research cohort. Our findings indicate that the majority of genetic variations, including those with apparent additive effects (i.e. significant GWAS loci), do not participate in pairwise interactions, or their interactions have negligible effects that cannot be detected even with a sample size of 3.6 million individuals. Furthermore, our results rule out the presence of epistatic interactions with effect sizes greater than 1.10mm for all allele pairs with frequencies down to 5%. While our result is inline with the previous findings showing mainly additive variance for height^2,5,12^, there are a few caveats worth discussing.

First, there are many plausible biological configurations under which two loci can interact. Biological epistasis can arise due to multiple factors. These include interactions at the molecular level, interactions at the regulatory levels such as transcription and translation, nuanced biochemical kinetic interactions like antigen recognition and substrate competitions, and broader functional redundancies within the pathways or interactions across pathways. The term “epistasis”, as defined by Fisher, refers to non-additivity, which is not an inherent biological condition but rather a modelling concept. In that sense, the lack of statistical evidence for pairwise interactions does not preclude the existence of gene networks that interact. This shows that our statistical model cannot capture any additional variance for height beyond marginal additive effects. This further confirms the robustness of the additive model in approximating height heritability.

Second, epistasis is only detectable when genetic variation is associated with phenotypic variance. For example, in a biallelic SNP, the mean trait value across the three genotypes may remain constant while the variance of trait values increases or decreases per allelic dosage. In this scenario, there may be no apparent additive effect, meaning there is no linear relationship between the trait mean values and allelic dosage. However, there is a significant epistatic effect, as the variance of the three genotypic groups is significantly different. This variance effect can be considered a signature of epistasis and has been previously studied in relation to other human traits and diseases^24–26^. Detection of genetic variation associated with phenotypic variance requires exceedingly large sample sizes because variance has a larger sampling error than the mean. It is quite likely that at our current cohort size we are underpowered to detect variance contributed by pairwise interaction independently of additive effects. Alternatively, it is entirely possible that we missed interactions between variants without apparent additive effects that do not reach genome-wide significance but they contribute to trait variance. Testing for those variants requires an exhaustive search of all pairwise interactions^27,28^ that is not addressed by our two-stage strategy.

Third, we restricted our analysis to only distal interactions that map greater than 2Mb from each other. The exclusion of proximal interactions (less than 2Mb apart) was primarily executed to circumvent the additive effect of untyped variants with low levels of LD colluding the test of epistasis^29,30^. While others used smaller (< 1Mb)^31^ or larger thresholds (< 5Mb)^26^ to protect against this effect, near-range interactions between haplotypes are reported^32^. Recent findings indicate that local epistasis in the form of haplotype interactions tends to be more circumstantial than global pairwise interactions in seven complex traits, including height^32^. In view of this, it is quite possible that by focusing on distal interactions, we overlooked the haplotype effects, where the majority of epistasis is concentrated.

Fourth, we only considered interactions between the common variants with MAF > 1%. While research in model organisms suggests that rare variants may disproportionately exhibit non-additive effects^33^, evidence of non-additivity among rare-variants in humans is mostly restricted to the recessive conditions^34^ and disease modifying interactions^35^. We speculate that we might have missed important pairwise interactions between rare variants and the common genetic background that could explain the observed preponderance of diminishing return epistasis (negative epistasis)^36,37^ among sub-significant interactions. Nevertheless, analysis of inbreeding depression in human height is inconsistent with significant epistatic effects from rare variants^38^.

Fifth, a notable limitation of our study is that our investigation of genetic interactions for height was confined exclusively to individuals of European descent. This focus restricts the generalizability of our findings, as genetic architectures and the distributions of variant effect sizes can vary significantly across different populations. Such variations can be due to differences in allele frequencies, environmental factors, and population-specific linkage disequilibrium structures, all of which can influence the detection and interpretation of genetic interactions. Consequently, while we report minimally meaningful evidence of genetic interactions influencing height within the European population, these results may not necessarily hold true for other groups. It remains an open question whether similar patterns of genetic interactions will be observed in non-European populations.

Overall, by analysing pairwise interactions in adult human height across 3.6 million individuals, we confirmed previously reported minimal contribution of epistasis in human height. This understanding provides a clear delineation of genetic influences on height and supports the focus on additive effects in genetic studies of complex traits. Future investigations into the genetic basis of height may benefit from concentrating on the identification and characterization of additive genetic variants rather than non-additive interactions, given their limited explanatory power observed in this study.

## Materials & Methods

### Study recruitment and data collection

The study involved the analysis of standing height measures among the research participants of 23andMe Inc., a personal genetics company. The study participants provided informed consent and took part in the research online, under a protocol that was approved by the external Association for the Accreditation of Human Research Protection Programs, an accredited institutional review board, and Ethical and Independent Review Services. Those who consented to participate were included in the analysis, based on the status of their consent at the time data analyses began. Participants were invited to complete web-based surveys, which collected information about their anthropometric traits including standing height. Reported height values were adjusted for age and top 5 PCs. To account for potential systematic differences in the distribution of the predictor variable, height, across sexes, we applied a quantile normalisation procedure to the height measurements of male and female participants separately. For each sex group (male and female), height measurements were first ranked. Subsequently, corresponding height values from each group were aligned to the same quantiles of the normal distribution, effectively standardising the scale and distribution of height across sexes.

### Genotyping, variant calling and imputation

Saliva samples were used for DNA extraction and genotyping using one of five genotyping platforms. The v1 and v2 platforms were based on the Illumina HumanHap550 BeadChip and contained approximately 560,000 SNPs, including about 25,000 custom SNPs selected by 23andMe Inc. The v3 platform was based on the Illumina OmniExpress BeadChip and contained approximately 950,000 SNPs and custom content to improve the overlap with our v2 array. The v4 platform was a fully custom array of around 950,000 SNPs and included a lower redundancy subset of v2 and v3 SNPs with additional coverage of lower-frequency coding variation. The v5 platform was based on the Illumina Global Screening Array, consisting of approximately 654,000 preselected SNPs and about 50,000 custom content variants.

Samples with genotyping call rates less than 98.5% were reanalyzed. Participants whose analyses failed repeatedly were recontacted by the 23andMe customer service to provide additional samples, as done for all 23andMe customers. After phasing genotyping data in SHAPEIT5, imputation was subsequently performed separately for each genotyping platform in Beagle 5.4 The participant genotype data were imputed against a compendium of imputation panels that combines the 23andMe multi-ancestry panel (including 14,403 samples from 23andMe augmented with publicly available panels, phase 3 1000 Genomes Project 2 and GTEx v8 3) with three independent publicly available reference panels including the publicly available Human Reference Consortium (HRC), and UK BioBank (UKBB) 200K Whole Exome Sequencing (WES).

Imputed genotypes were subjected to quality control processing that involved masking variants with significant HWE deviation (p < 1e-20), low allele-frequency (MAF < 0.1%), low-genotyping call rate (<90% across all individuals), low-imputation quality (average r^2^ < 0.5 across batches or minimum r^2^ < 0.3), variants with significant batch effect (analysis of variance (ANOVA) F-test across batches, p < 1e-50), variants with large sex-effect (via ANOVA of SNP-dosage, removing variants with sex R^2^ > 0.1) and variants that fail Mendelian transmission test (p < 1e-20). Following quality control processing, the final imputed dataset contained 99.7 million variants per individual.

### Genome-wide association analysis (GWAS)

The 23andMe genetic ancestry classification algorithm was used to select participants of European ancestry. To ensure a maximal set of unrelated individuals, a segmental identity-by-descent (IBD) estimation algorithm was used. Individuals were considered related if they shared more than 700 cM IBD, which includes regions where both genomic segments on homologous chromosomes or one of the genomic segments were shared. This level of relatedness is equivalent to the minimum expected sharing between first cousins in an outbred population.

A total of 63,528 high-quality autosomal variants present in all five genotyping platforms were used to perform a principal component analysis. The analysis was performed on a randomly sampled subset (513K) of European participants from all the genotyping platforms. PC scores for participants not included in the analysis were obtained by combining the eigenvectors of the analysis and the variant weights and projecting them.

Linear regression was used to perform association analysis between genotyped and imputed variant dosage data of quantile normalised height, assuming an additive model of allelic effects. For each GWAS analysis, age, sex, the top five principal components, and the genotyping platforms were included as covariates in the regression model. The p-value for the association test was calculated using a likelihood ratio test comparing the goodness-of-fit of the null model (i.e. height ∼ covariates) versus the main model with variant effect (i.e. height ∼ *D*_snp_ + covariates, where *D*_snp_ ∈ {0, 1, 2}). Association statistics were derived for both autosomal and the X-chromosome variants, treating males as homozygous diploid for the observed variant. The LD-score regression method was applied to investigate polygenicity and correct for the residual population stratification and derive the narrow-sense heritability. The credible set for individual GWAS hits (p< 5e-8) was calculated using The Wellcome Trust Case Control Consortium method^29,30^. Variant coordinates are reported according to the GRCh38 assembly and annotated in VEP v109.2 using dbSNP v153, GENCODE v43.

We identified the genomic boundaries of each additive association, by considering all variants in the vicinity of a genome-wide significance lead variant with p< 1e-6. If two loci were separated by less than 250 kb, we grouped them into one locus. Within each locus, we identified the index variant as the one with the smallest p-value. We then annotated functional variants that were within 500Kb of the locus and in LD with the lead variant (r2 > 0.5). We set the minimum distance between loci as 250Kb, grouping closer hits into one locus.

### Translating effect sizes from quantile-normalised height to millimetres

The quantile normalisation process transforms the phenotype to a standard normal distribution, resulting in effect sizes expressed in terms of standard deviations (SD) of the normalised data. For interpretability, these effect sizes are converted back to millimetres using the standard deviation of the original height distribution. Briefly, the standard deviation of the original height measurements (*σ*) in mm(s) was calculated from the raw height data. The effect sizes from the GWAS (β_*SD*_), expressed in terms of standard deviations of the quantile-normalised height, were converted to millimetres using the following formula:

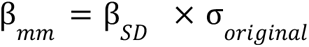

Where, β*_mm_* represents the effect size in millimetres, β*_SD_* is the effect size in standard deviations of the quantile-normalised height and σ*_original_* is 73.75 millimetres, as estimated from our data.

### Pairwise SNP-interaction discovery (GWAS SNP-SNP interaction model)

To test for G×G interactions, we employed a hierarchical modelling approach that incrementally integrates genetic information. This methodological framework is designed to identify and quantify the contributions of individual SNPs and their interactions to phenotypic variance in height. The analysis was structured around three models:

1. **Null Model (H_0_):** The baseline model posits that height is influenced solely by the additive effect of a single SNP (SNP1), along with a set of covariates. The model is formally described as:

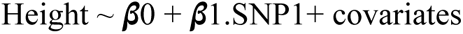

This model serves as a reference point, allowing us to assess the contribution of a single SNP to height variation.
2. **Additive Model (H_1_):** Expanding upon the null model, this model incorporates an additional SNP (SNP2) to evaluate its additive effect alongside SNP1. The model is represented as:

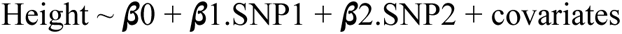

Here, the focus shifts to understanding how the combined additive effects of two SNPs contribute to height.
3. **Interaction Model (H_2_):** This model introduces an interaction term between SNP1 and SNP2 to the additive model, aiming to capture the joint effect of these SNPs on height. The formulation of this model is:

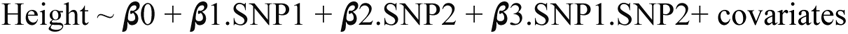

The inclusion of the interaction term (*β*3) allows for the investigation of whether and how the interaction between SNP1 and SNP2 influences height beyond their individual additive effects. Covariates included in all models encompass age, sex, the top five principal components (PCs) to adjust for population stratification, and genotyping platform variables. The significance of the SNP-SNP interactions was assessed using a likelihood ratio test, comparing the goodness-of-fit between the interaction model (H_2_) and the simpler models (H_1_ and H_0_), with a null hypothesis that *β*3 = 0. Overall we tested 564,453 SNP-pairs. We adopted the Bonferroni-corrected significance threshold of (0.05/564,453 = 8.90e-8) as the universal threshold for calling significant pairwise interactions.

To ensure a balanced distribution of samples when stratifying on genotypes, we restricted the interaction analysis to common variants (MAF > 1%). We further removed any interactions with a proximal distance of less than 2Mb. This ensures that the identified interactions are not due to the “phantom epistasis”^30^ effect arising from the LD structure of the locus.

### Fully saturated model with additive and non-additive components

We explore the complexities of pairwise genetic interactions and their contributions to height variance in a generalised linear framework. Recognizing that dominance effects within gene pairs might obscure or falsely suggest SNP interactions, we adopted a nested model approach. This methodological framework allows us to discern whether SNP interactions exert a significant influence on height beyond the dominance effects of the interacting alleles. To achieve this, we constructed and compared two linear models:

1. **Null Hypothesis Model (H_0_) - Base Model with Dominant Terms:** This model incorporates both the additive and dominance effects of two single nucleotide polymorphisms (SNPs), SNP1 and SNP2, as well as a set of covariates. The model is specified as follows:

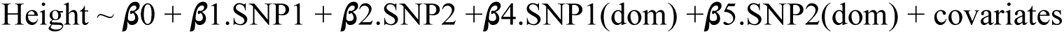

This setup models height as a function of the additive effects of SNP1 and SNP2, their respective dominance deviations, and additional covariates.

1. **Alternative Hypothesis Model (H_1_) - Fully Partitioned Model (Full Model):** Expanding upon the base model, this formulation introduces interaction terms to evaluate the joint effects of SNP1 and SNP2, including their interactions with each other’s dominance deviations:

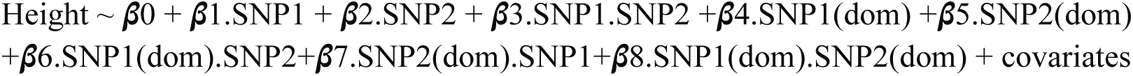

This comprehensive model assesses the significance of epistatic interactions including additive-by-additive, additive-by-dominance and dominance-by-dominance between pairs of SNPs after accounting for their dominance effects and includes the same covariates as the base model. For both models, the additive genetic effect is coded numerically (0, 1, 2) corresponding to non-carrier homozygotes, heterozygotes, and effect-allele carrier homozygotes, respectively. Dominance deviation is coded as 1 for homozygous carriers of the effect allele and 0 otherwise. Covariates in these models encompass sex, age, interactions between sex and age, principal components (pc.1 to pc.5) to adjust for population stratification, interactions of the first two principal components with sex and age, and interactions between each SNP with age and sex, providing a comprehensive control for potential confounders.

Model comparison and the significance of SNP interactions were evaluated using a likelihood ratio test, contrasting the full model against the base model. This statistical approach allows us to determine whether the inclusion of interaction terms significantly improves the model fit, indicating a substantive contribution of SNP interactions to the phenotypic variance observed in human height.

In order to scientifically determine whether the interaction effect sizes of SNP pairs were consistently smaller than their additive effects, we employed a one-sided signed binomial test. Specifically, for each SNP pair, we tested the null hypothesis that the interaction effect size was equal to or greater than the additive effect size, against the alternative hypothesis that the interaction effect was smaller.

### GxG power estimation

We calculated the power to detect genetic interactions between two loci across a population sample of 3.6 million individuals. We assume that the estimated effect size under the null hypothesis (i.e. no interaction) is distributed as N(0, 1). Under the alternative hypothesis (interaction effect of size *β*), we assume that the estimated effect size is distributed as N(*β*, *se*^2^). We simulated height data assuming that the distribution is N(*μ*, *σ*^2^), where *μ* = 1723.15mm and *σ*^2^=(73.75mm)^2^, based on the observed height distribution in our cohort. We simulate genetic data by drawing independent samples from the binomial distribution *xi*∼*Bin*(*N* = 2, *f*1) and *ui*∼*Bin*(*N* = 2, *f*1), where *f*1 and *f*2 are the minor allele frequencies (MAFs) for the two loci. In our simulation, we consider a range of MAFs for *f*1 (0.01 ≤ *f*1 ≤ 0.5) and fix *f*2 at 0.5, representing SNP 1 and SNP 2, respectively. We then fit a linear model (height ∼ *μ* + β_1_*x_i_* + β_2_*u_i_* + β_3_*x_i_u_i_*+ ε) to obtain an estimate of *se* under the null. Given the above, we define the test statistic as chi-square distributed with 1 degree of freedom. The non-centrality parameter (NCP) is given by: *NCP*= (β_3_/*SE*_β_3__)^2^ Using the NCP and a χ^2^ test, we estimated the statistical power of identifying GxG interactions at a p-value threshold of 8.9e-8. The power was calculated across a range of allele frequencies for SNP 1 (0.01 ≤ *f*1 ≤ 0.5).

## Acknowledgements

We would like to thank the research participants and employees of 23andMe Inc. for making this work possible.The following members of the 23andMe Research Team contributed to this study:

Stella Aslibekyan, Adam Auton, Elizabeth Babalola, Robert K. Bell, Jessica Bielenberg, Ninad S. Chaudhary, Zayn Cochinwala, Sayantan Das, Emily DelloRusso, Payam Dibaeinia, Sarah L. Elson, Nicholas Eriksson, Chris Eijsbouts, Pierre Fontanillas, Davide Foletti, Will Freyman, Julie M. Granka, Chris German, Éadaoin Harney, Alejandro Hernandez, Barry Hicks, David A. Hinds, Michael V Holmes, M. Reza Jabalameli, Ethan M. Jewett, Yunxuan Jiang, Sotiris Karagounis, Matt Kmiecik, Katelyn Kukar, Alan Kwong, Keng-Han Lin, Yanyu Liang, Bianca A. Llamas, Aly Khan, Steven J. Micheletti, Matthew H. McIntyre, Meghan E. Moreno, Priyanka Nandakumar, Dominique T. Nguyen, Steve Pitts, G. David Poznik, Alexandra Reynoso, Shubham Saini, Morgan Schumacher, Leah Selcer, Anjali J. Shastri, Jingchunzi Shi, Suyash Shringarpure, Keaton Stagaman, Teague Sterling, Qiaojuan Jane Su, Joyce Y. Tung, Susana A. Tat, Vinh Tran, Xin Wang, Wei Wang, Catherine H. Weldon, Amy L. Williams, Peter Wilton.

## Data availability

The full GWAS summary statistics for the 23andMe discovery data set will be made available through 23andMe to qualified researchers under an agreement with 23andMe that protects the privacy of the 23andMe participants. Datasets will be made available at no cost for academic use. Please visit https://research.23andme.com/collaborate/#dataset-access/ for more information and to apply to access the data.

