## Supplementary Materials for "Analysis of 3.6 million individuals yields minimal evidence of pairwise genetic interactions for height"

### Supplementary Notes

#### Analysis of dominance across the 69 variants tested in the fully saturated model

We identified five variants with significant dominance effect at our lenient threshold ($p < 1e-4$) (**Supplementary Table 5**). Consistent with previous findings[1], we observed that the largest contribution to dominance deviation occurs among alleles that are considerably common with MAF ranging from 27% to 40% in our cohort. The top hit, rs11205303 (p-value= 2e-16), is a missense variant in *MTMR11*, which encodes a member of the myotubularin tyrosine phosphatase family. Unlike other classes of myotubularins, MTMR11 does not interact with themselves or other myotubularins[2]. The rs1926872 lead variant, is an intergenic variant that maps between *COLGALT2* and *TSEN15* and is an eQTL for both genes. The rs7832643 variant is located in the intronic region of *PLEC*, which encodes plectin, a relatively large protein (~ 500 kDa) involved in the cytoskeleton integrity. The rs888922 hit maps to the intronic region of *GFPT2* and is an eQTL for this gene. *GFPT2* encodes for the enzyme Glutamine--fructose-6-phosphate transaminase 2 that has a central role in glycosylation. The rs3843751 variant is an intronic variant mapping to the intronic region of *SLC44A2* which encodes the solute carrier family 44, member 2 protein. This protein is part of the solute carrier (SLC) group, which is a diverse family of membrane-bound transporters responsible for moving a variety of substances across cellular membranes. The last hit, rs2100374, lacks coding or eQTL variant-to-gene resolution and maps to the intergenic region between *ELOVL5* and *GCLC*. Neither of these loci were previously reported in the dominance analysis of standing height in the UK Biobank[1].

#### Variance explained by non-genetic components in the saturated model

In assessing the contribution of covariates and their interactions to the total height variance (**Supplementary Figure 9**) we identified that age is a substantial contributor to variance, explaining 15.58% (IQR range: 10.68- 20.02%, SE=0.80) of the variance in the model, making it the most impactful single covariate. Sex, while included as a factor, contributes less to the model, with a notably smaller percentage of variance explained (0.38%, SE=0.35). Among the principal components, PC3 stands out by explaining a significant percentage of variance (4.40%, SE=0.20) , second only to age. This suggests that PC3 captures key variation related to the phenotype, which may not be directly linked to age or sex but represents other underlying biological or environmental gradients. The remaining PCs (PC1, PC2, PC4, and PC5) each account for a notably smaller fraction of the variance, suggesting that the factors they represent have a lesser impact on the phenotype (**Supplementary Figure 9**).

Interaction terms between non-genetic covariates bring additional insights. The interaction between age and PCs, along with the interaction between sex and age, contributes significantly to model variance. Quantitatively, on average, the interaction of age by PC1 and PC2 explains 0.005% and 0.01% of the total phenotypic variance, and 8.9 % and 21.6% of the total variance explained by the model. This suggests that the effect of age on height is modulated by both sex and the top two PCs. Conversely, other interactions involving sex, such as sex by PC1 and sex by PC2, have less influence on phenotypic variance (**Supplementary Figure 10**). These findings suggest a complex interplay between genetic factors represented by principal components and non-genetic factors such as age and sex in influencing height variance.

### Supplementary Figures

**Supplementary Figure 1:** Manhattan plot and MAF stratified QQ-plot for adult height GWAS.

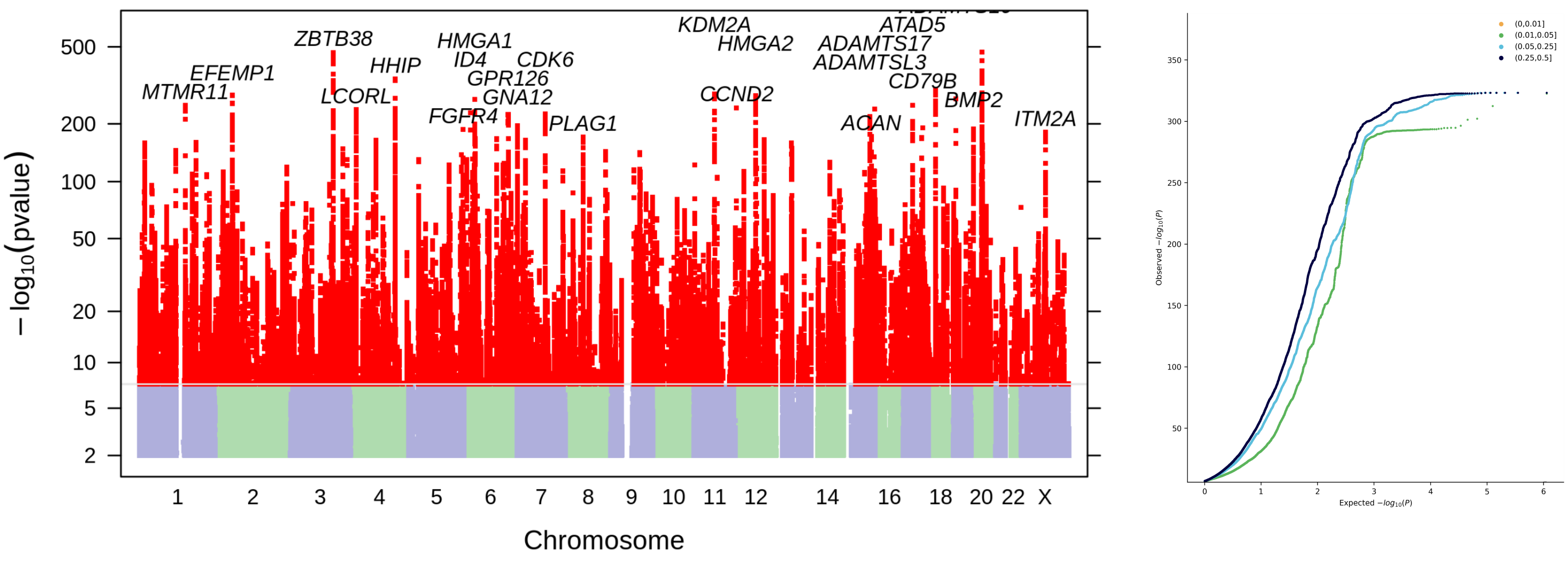

**Supplementary Figure 2:** Relationship between minor-allele frequency and estimated additive effect sizes of effect alleles. Each dot represents one of the 1,063 independent GWAS hits.

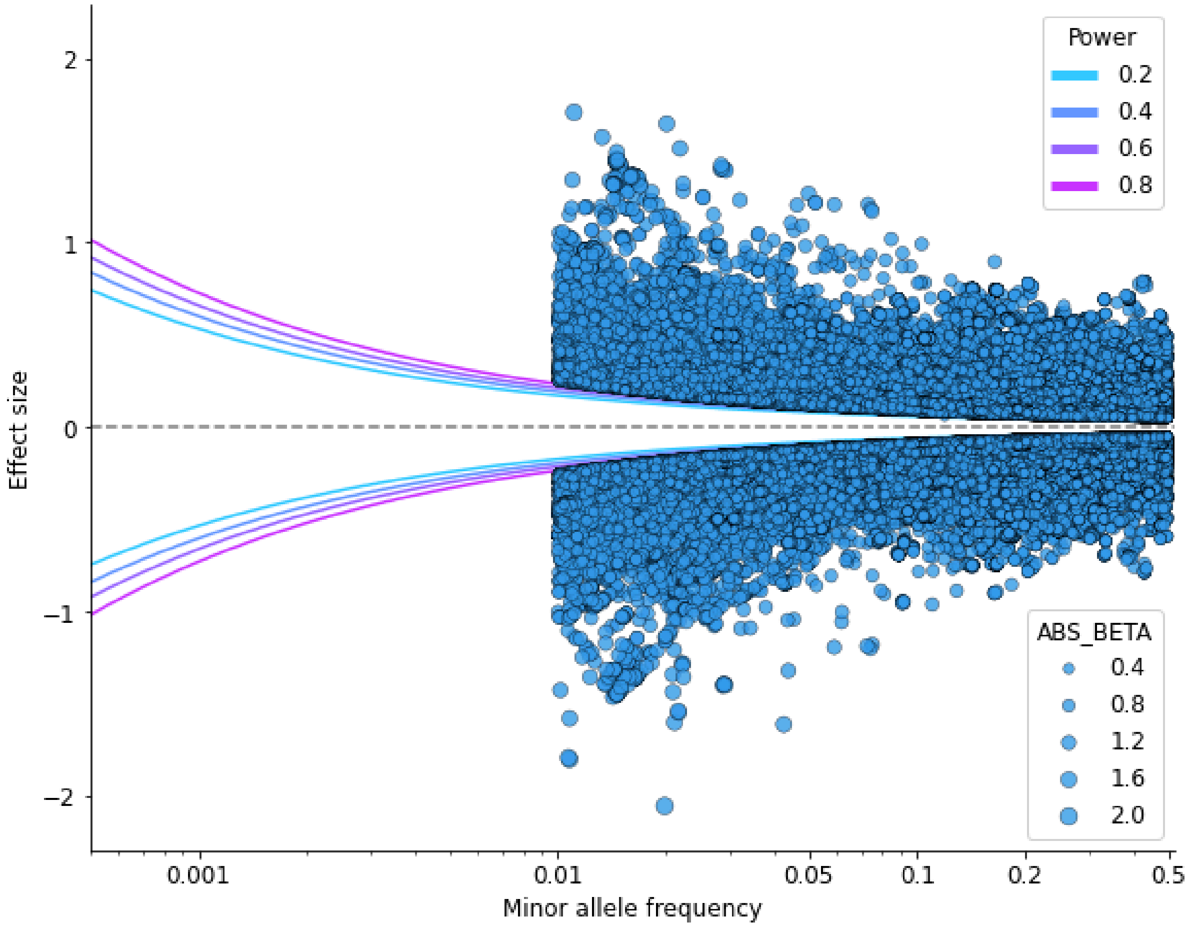

For presentation purposes, the SNP effect estimates on the y-axis are scaled by a factor of ten for clearer visualisation and expressed in quantile normalised standard deviation (s.d.) per effect allele. We show four curves representing the theoretical relationship between frequency and expected magnitude of variant effect detectable at genome-wide significant threshold ($p < 5e\text{-}8$) with a statistical power of 20%, 40%, 60% and 80%.

**Supplementary Figure 3:** Histogram of the pairwise distance between independent SNP associations on chromosome 1-22 and the X chromosome.

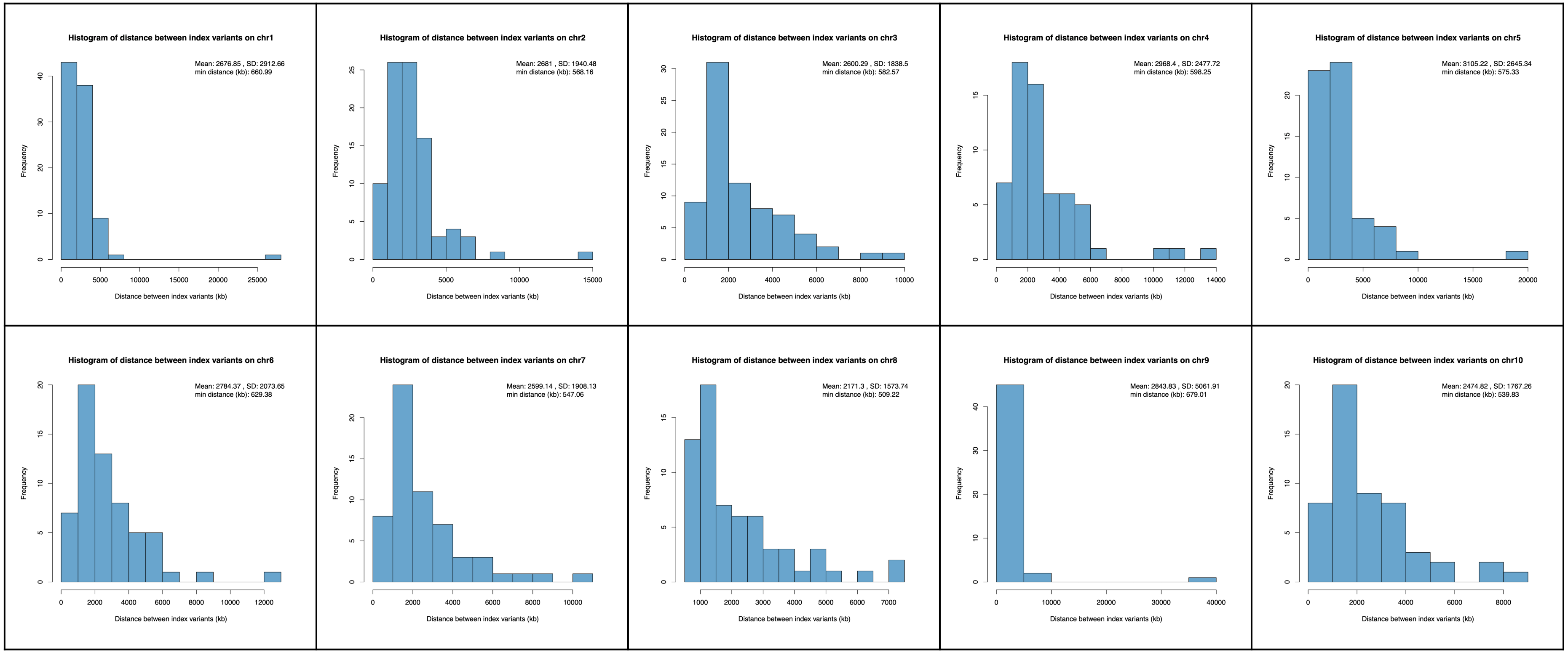

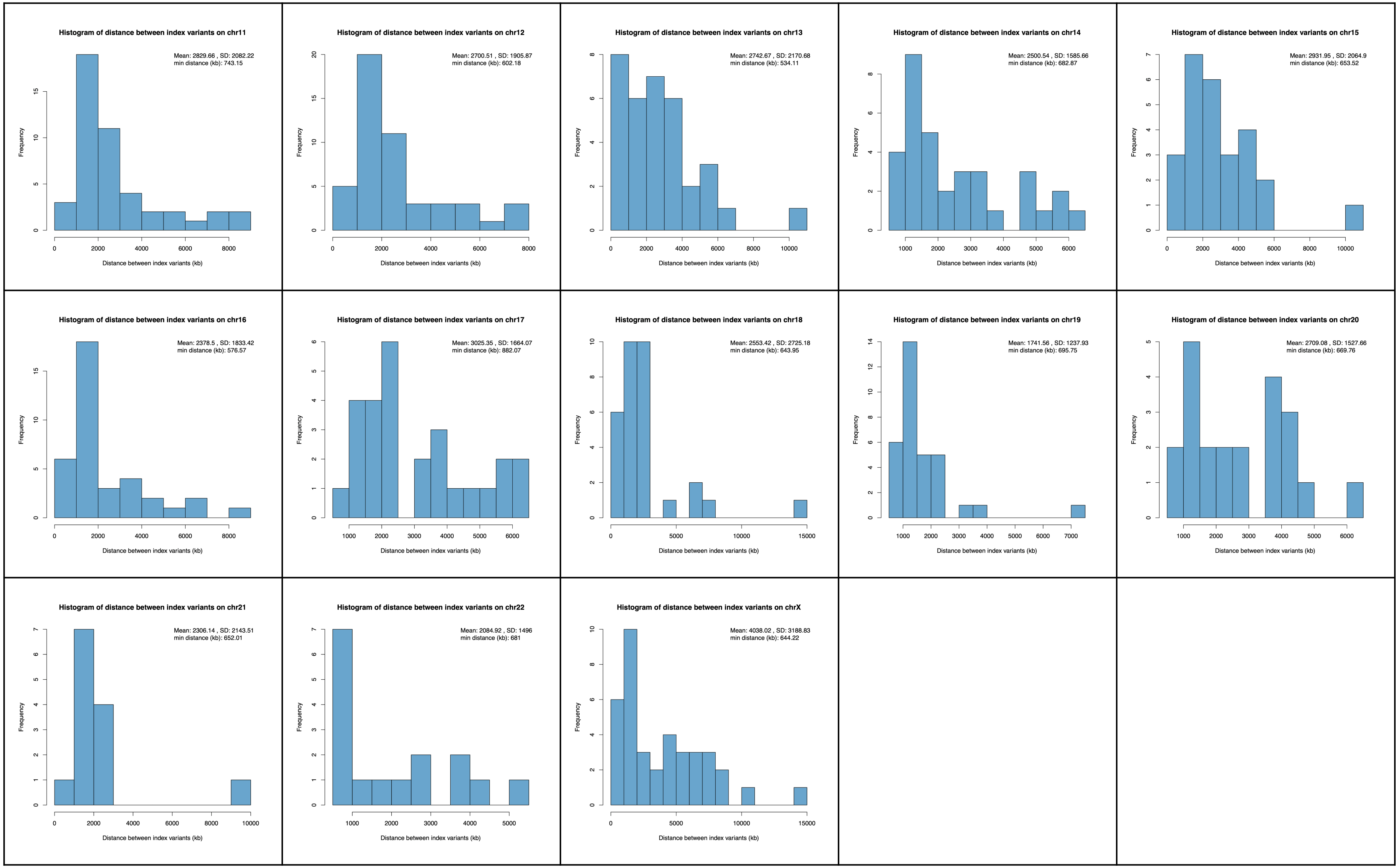
Histogram of distance between index variants on chr11
Mean: 2829.66 , SD: 2082.22
min distance (kb): 743.15
Histogram of distance between index variants on chr12
Mean: 2700.51 , SD: 1905.87
min distance (kb): 602.18
Histogram of distance between index variants on chr13
Mean: 2742.67 , SD: 2170.68
min distance (kb): 534.11
Histogram of distance between index variants on chr14
Mean: 2500.54 , SD: 1585.66
min distance (kb): 682.87
Histogram of distance between index variants on chr15
Mean: 2931.95 , SD: 2064.9
min distance (kb): 653.52
Histogram of distance between index variants on chr16
Mean: 2378.5 , SD: 1833.42
min distance (kb): 576.57
Histogram of distance between index variants on chr17
Mean: 3025.35 , SD: 1664.07
min distance (kb): 882.07
Histogram of distance between index variants on chr18
Mean: 2553.42 , SD: 2725.18
min distance (kb): 643.95
Histogram of distance between index variants on chr19
Mean: 1741.56 , SD: 1237.93
min distance (kb): 695.75
Histogram of distance between index variants on chr20
Mean: 2709.08 , SD: 1527.66
min distance (kb): 669.76
Histogram of distance between index variants on chr21
Mean: 2306.14 , SD: 2143.51
min distance (kb): 652.01
Histogram of distance between index variants on chr22
Mean: 2084.92 , SD: 1496
min distance (kb): 681
Histogram of distance between index variants on chrX
Mean: 4038.02 , SD: 3188.83
min distance (kb): 644.22
Frequency
Distance between index variants (kb)

The panel shows histograms of the distances between independent SNP associations on chromosomes 1-22 and chromosome X. The histogram displays the distribution of distances (in kilobases, kb) between index variants. The frequency of distances is shown on the y-axis, while the distance between index variants is represented on the x-axis.

**Supplementary Figure 4:** P-value distribution and Q-Q plot of additive-by-additive genetic interactions from GWAS SNP-SNP interaction screen.

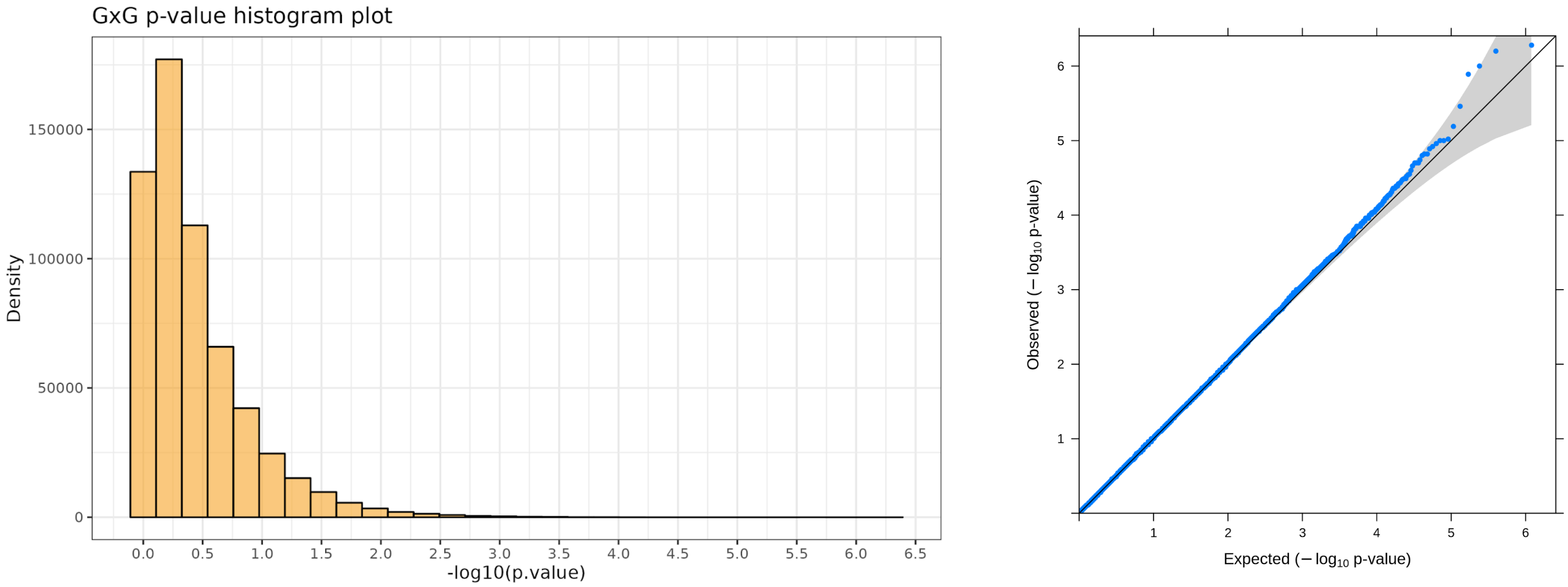

**Figure 4.a: Histogram of Discovery G×G P-values:** The histogram illustrates the distribution of interaction p-values derived from a genome-wide SNP-SNP interaction screen for height. The x-axis represents the negative logarithm to the base 10 of the G×G p-values, allowing for a more discernible visualisation of the distribution of small p-values. The y-axis denotes the frequency of each p-value range. **Figure 4.b: QQ Plot of Interaction P-values:** The QQ plot assesses the distribution of the observed interaction p-values against the expected distribution under the null hypothesis of no interaction. The x-axis plots the expected -log10 p-values if the data were perfectly following the expected distribution, while the y-axis plots the observed -log10 p-values from the GWAS SNP-SNP interaction data. Points falling along the diagonal line indicate adherence to the expected null distribution, while deviations from this line suggest an enrichment of significant p-values. The shaded area represents the 95% confidence interval under the null hypothesis, providing a visual cue to assess deviations that might indicate true associations.

**Supplementary Figure 5:** Sensitivity of interaction signal retention relative to p-value cutoff thresholds.

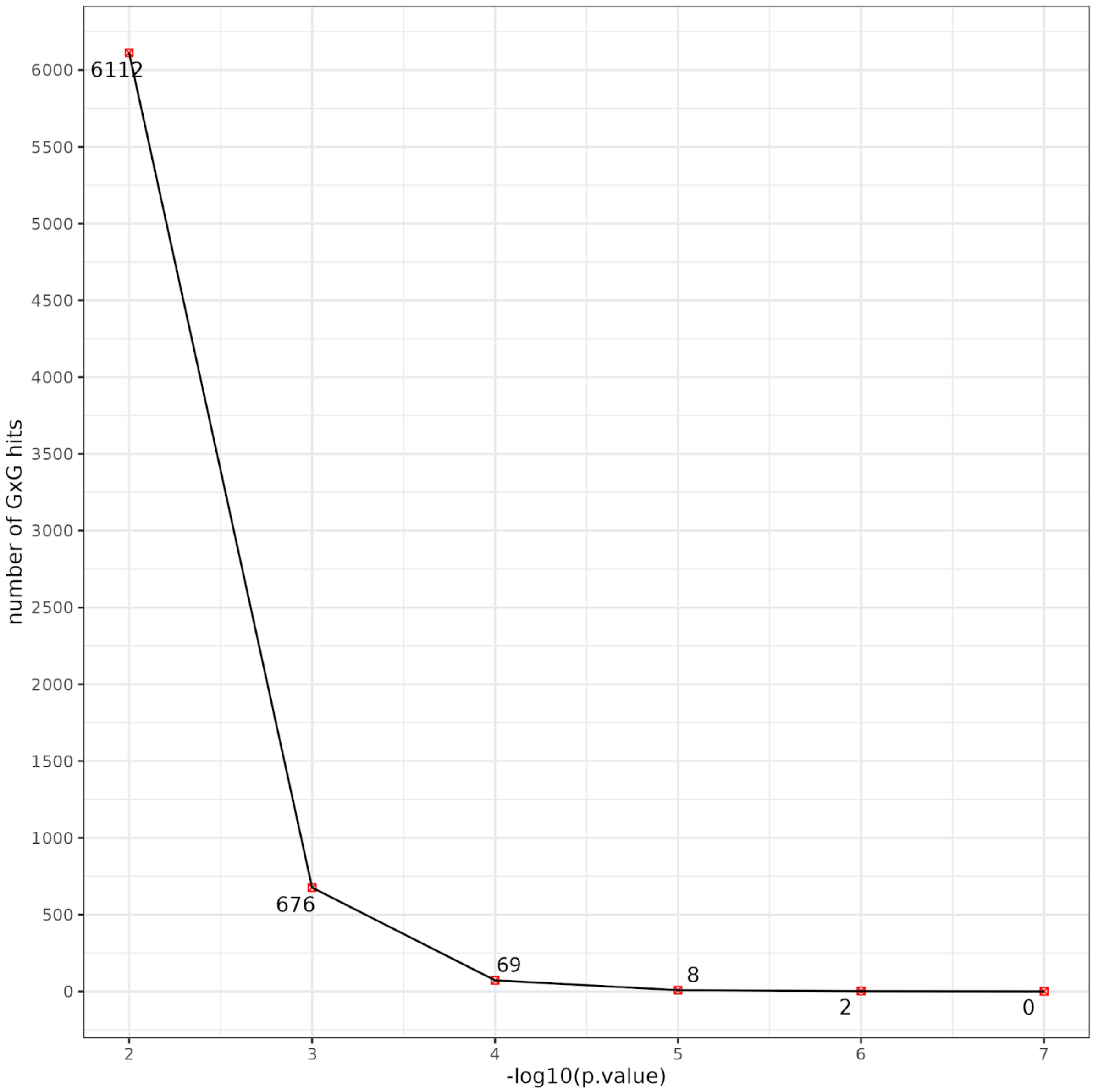

The line plot illustrates the relationship between various p-value cutoff thresholds (x-axis) and the number of interaction signals that remain above these thresholds (y-axis). The x-axis represents the -10log (G×G p-value), showcasing how stricter cutoffs (higher values on the x-axis) correlate with fewer retained interactions (lower values on the y-axis. This plot demonstrates the sensitivity of interaction signal detection to the choice of p-value threshold, emphasising the trade-off between signal strength and the cutoff adopted for controlling Type I error.

**Supplementary Figure 6:** Intrachromosomal interactions reaching empirical significance threshold (p < 1e-04).

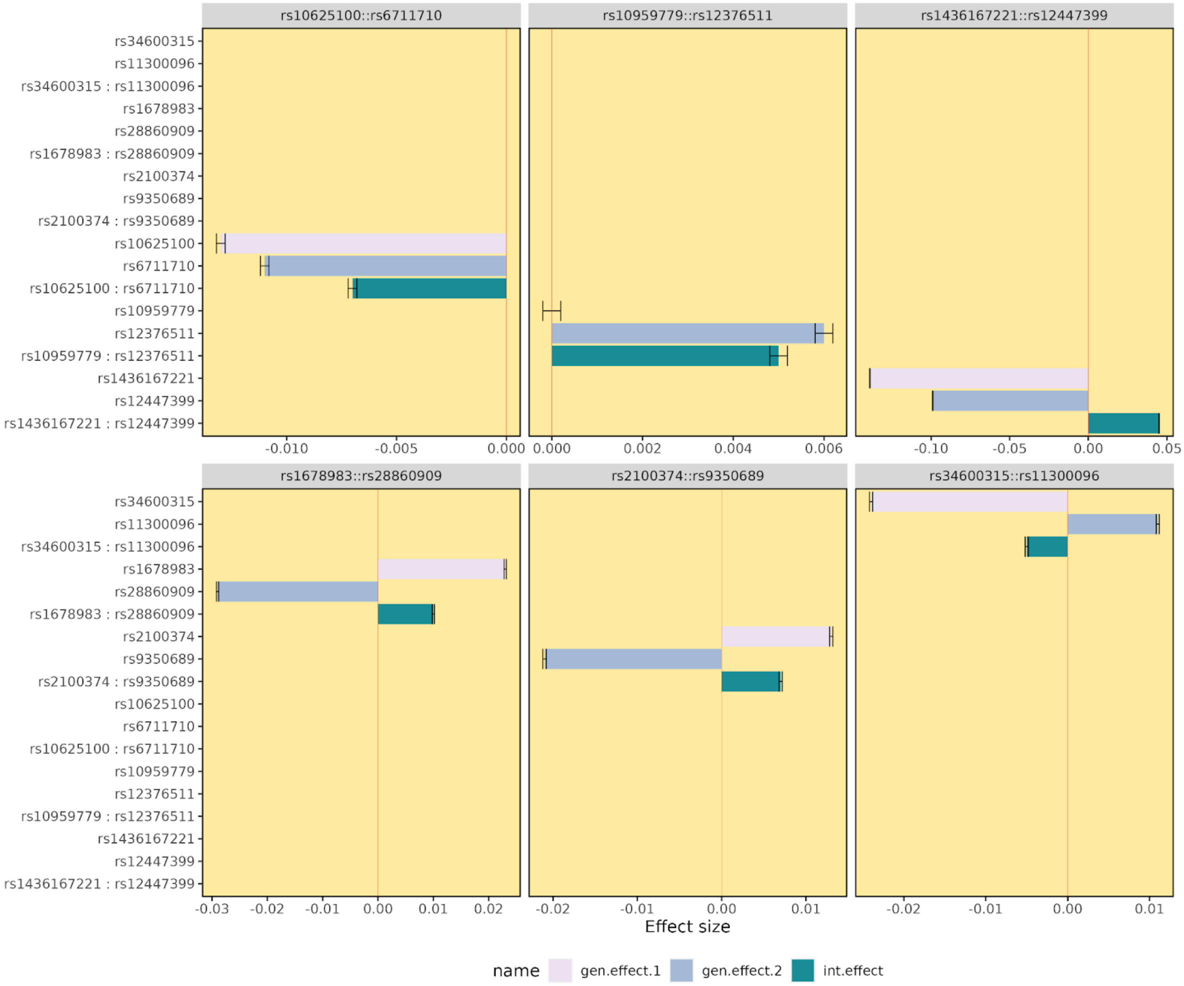

This bar plot visualises the effect sizes of significant cis-interactions (i.e. intrachromosomal interactions) between pairs of SNPs that have surpassed an empirical significance threshold (p < 1e-04). Each panel corresponds to a unique SNP pair, with the SNP identifiers listed along the y-axis. Within each panel, two coloured bars represent the additive effects of each SNP involved in the interaction: the first bar in pink indicates the additive effect of SNP1, and the second bar in blue denotes the additive effect of SNP2. The interaction effect of the SNP pair is represented by a bar in green. Error bars extending from each coloured bar capture the standard error associated with the effect size estimate. The x-axis is scaled to show the magnitude of the effect sizes for each type of genetic effect. The layout of the plot is designed to offer a comparative view of the strength of the individual SNP effects (expressed in standard deviations (s.d.) per minor allele) against their combined interaction effect, facilitating an interpretation of how these SNPs might influence a trait synergistically beyond their individual contributions.

**Supplementary Figure 7:** Percentage of phenotypic variance explained by additive, dominance and epistatic gene actions.

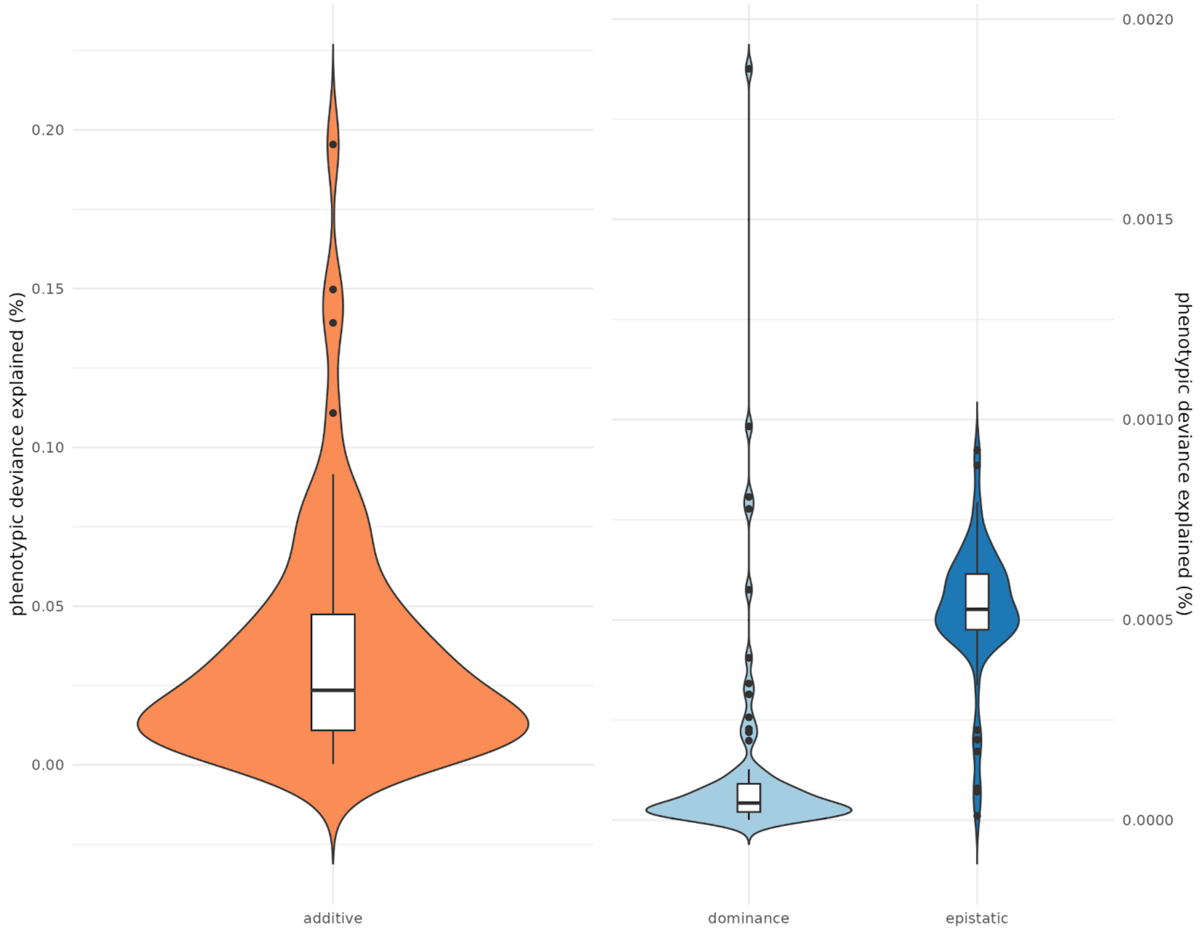

The violin plot illustrates the distribution of the percentage of phenotypic variance explained by additive, dominance, and epistatic (sum of additive by additive, additive by dominance and dominance by dominance) gene actions. Each violin represents the density distribution of the variance explained, with its width corresponding to the frequency of observations. Within each violin, a box plot is embedded, indicating the interquartile range (IQR) with a line denoting the median. Outliers are represented by dots outside the violins. The left violin, in orange, illustrates the additive gene action with a considerably wider distribution, suggesting a larger variation in the percentage of variance explained by this component. The right two violins, in blue, represent dominance and epistatic gene actions, respectively, and are plotted against a different y-axis scale due to their smaller range of variance explained. This separate scaling is necessary to visually accommodate the lower percentages of phenotypic variance explained by dominance and epistatic actions compared to the additive component. The right scale shows that the dominance and epistatic components contribute much less to phenotypic variance than the additive effects, with epistatic interactions showing the smallest distribution, indicative of its minor role in the variance explained. The figure highlights the varying contributions of genetic actions to phenotypic variance in height, with additive gene action being the most significant contributor, while dominance and epistatic actions have a notably smaller impact.

**Supplementary Figure 8.a:** Distribution of additive variance explained across 69 SNP pairs with suggestive evidence of interaction.

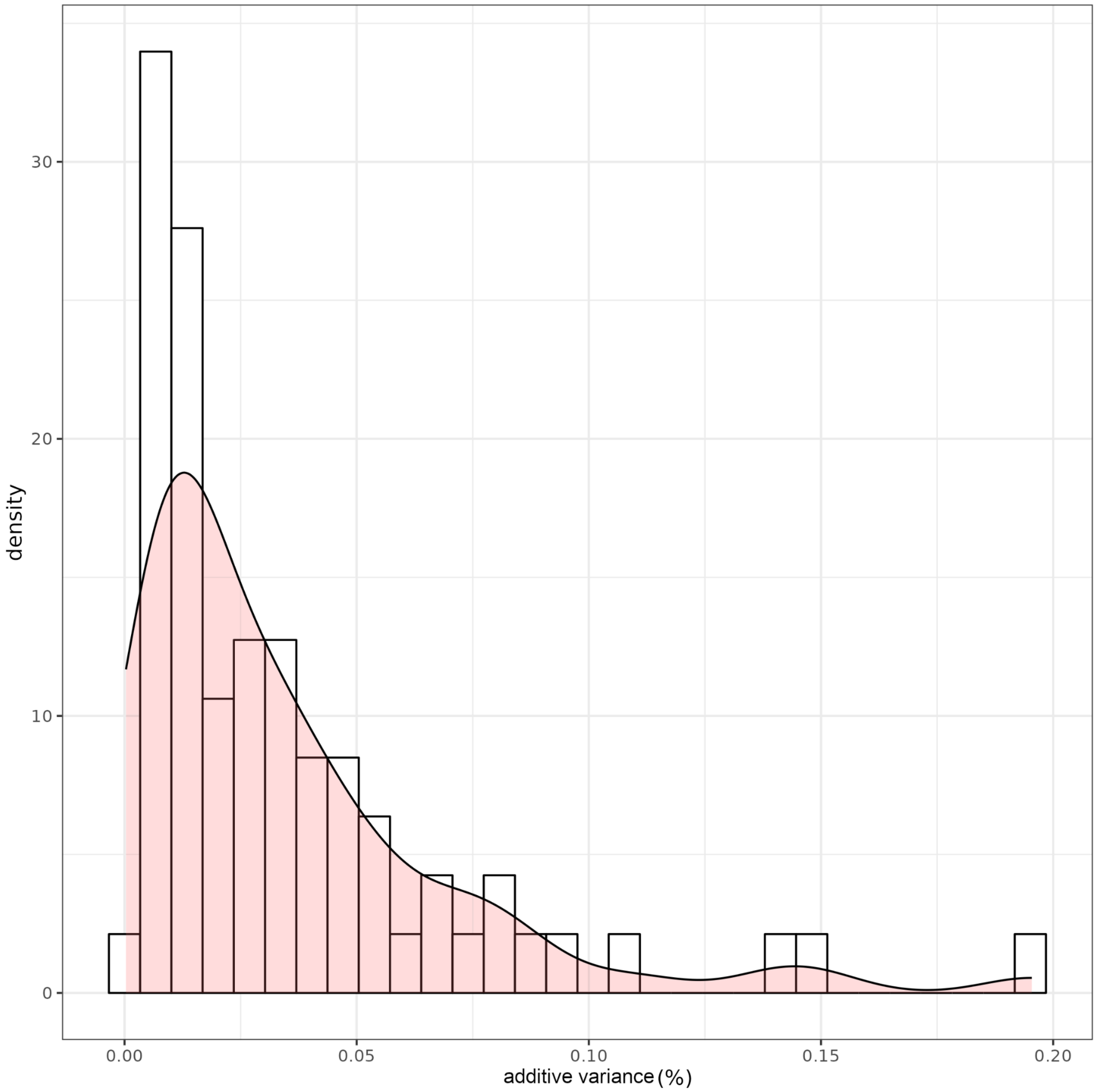

The histogram with an overlaid density plot illustrates the distribution of the percentage of additive variance explained for height. The histogram bars represent the frequency of data points within specific intervals of additive variance percentages, offering a visual representation of the distribution's shape and spread. The smooth curve represents the density plot, which estimates the probability distribution of the data. The plot shows the distribution's skewness toward lower percentages, indicating that most of the additive variance values are clustered near smaller values. This suggests that the additive genetic contribution to the phenotypic variance of height is generally modest across the dataset. No data points are observed beyond 0.20% additive variance, reflecting the limited contribution of individual additive effects to the overall phenotypic variance in this particular study.

**Supplementary Figure 8.b:** Distribution of dominance variance explained across 69 SNP pairs with suggestive evidence of interaction.

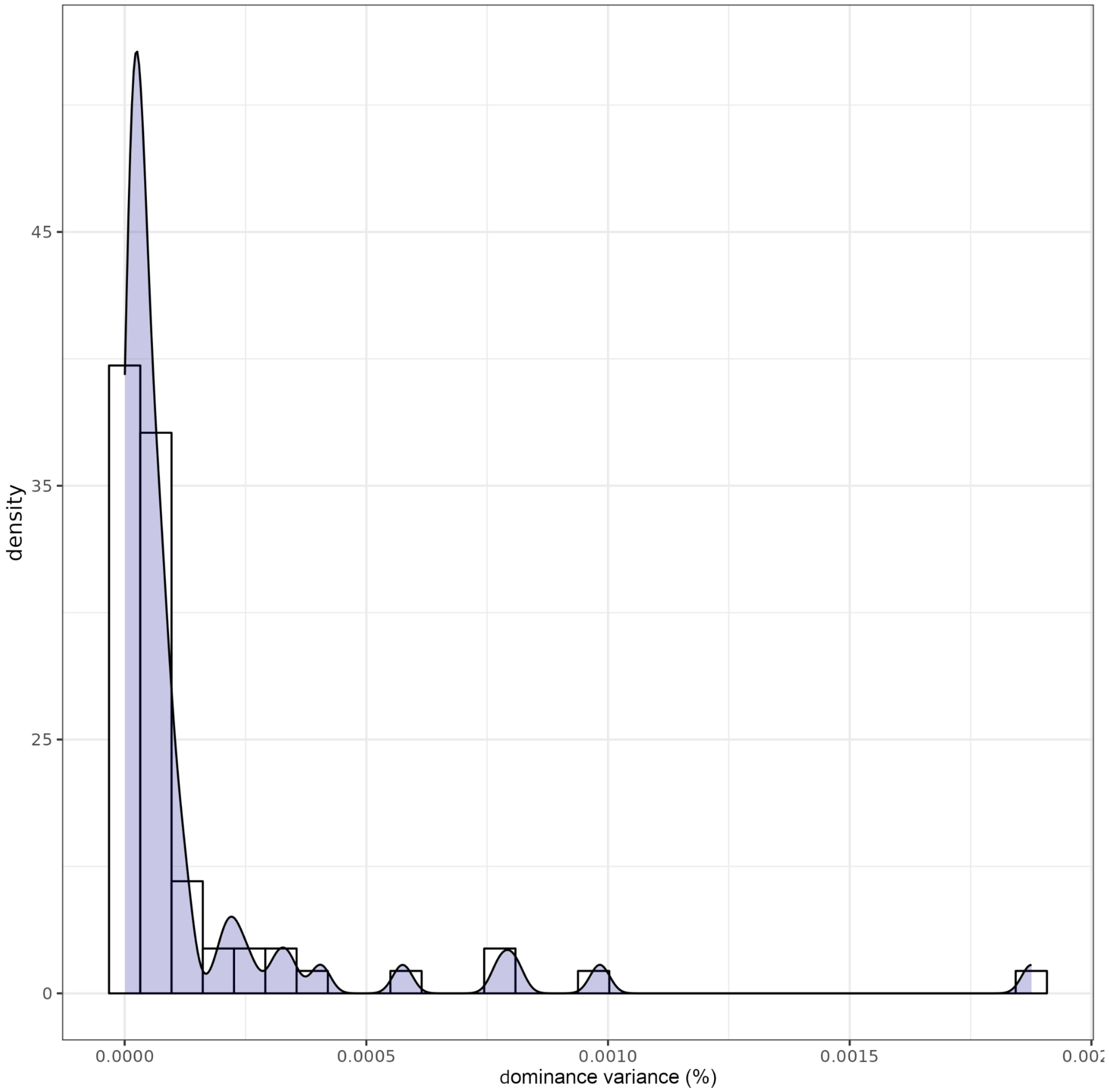

The histogram with an overlaid density plot illustrates the distribution of the percentage of phenotypic variance in height explained by dominance genetic effects. The x-axis represents the range of dominance variance percentages, and the y-axis denotes the density. The histogram bars indicate the frequency of dominance variance percentages observed, illustrating a distribution that is markedly skewed towards the lower end of the scale, with a long tail extending to the right. This skewness suggests that dominance effects are generally small, with most observed values close to zero and only a few reaching higher percentages. The superimposed black line represents the kernel density estimate, providing a smoothed approximation of the distribution shape. The distribution indicates that dominance genetic effects contribute a relatively minor proportion to the phenotypic variance of height in this population, with the vast majority of these effects explaining a very small amount of the variance. The presence of the long tail indicates occasional larger contributions, but these are clearly outliers within the dataset.

**Supplementary Figure 8.c:** Distribution of additive by additive (G×G) variance explained across 69 SNP pairs with suggestive evidence of interaction.

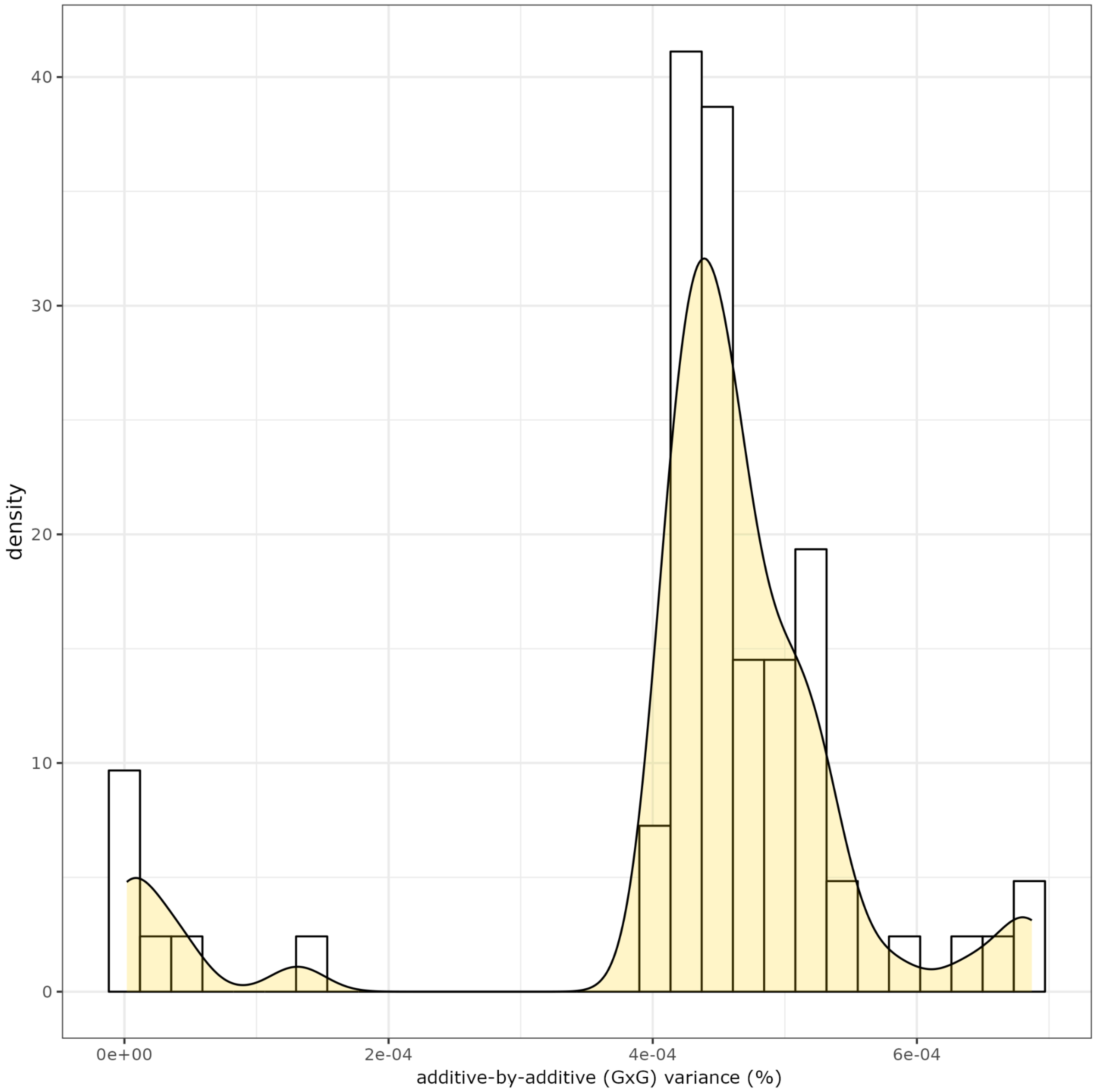

The histogram with an overlaid density plot illustrates the distribution of the percentage of phenotypic variance explained by additive-by-additive (G×G) interactions for height. The x-axis represents the range of G×G variance percentages, while the y-axis shows the density of these percentages within the dataset.The distribution is heavily skewed towards the lower end of the percentage scale, which indicates that the vast majority of additive by additive interactions explain a very small fraction of the phenotypic variance in height. The tail of the distribution extends to higher percentages, albeit these represent a minority of the cases. A smooth curve traces the density estimate across the distribution, providing a continuous representation of the underlying data structure. This plot emphasises the rarity of substantial additive-by-additive contributions to the overall variance in height, underscoring the relatively minor role that additive-by-additive interactions play in the genetic architecture of this phenotype within the studied population.

**Supplementary Figure 9:** Comparison of various non-genetic covariates' contribution to height variance, as explained in the fully partitioned model.

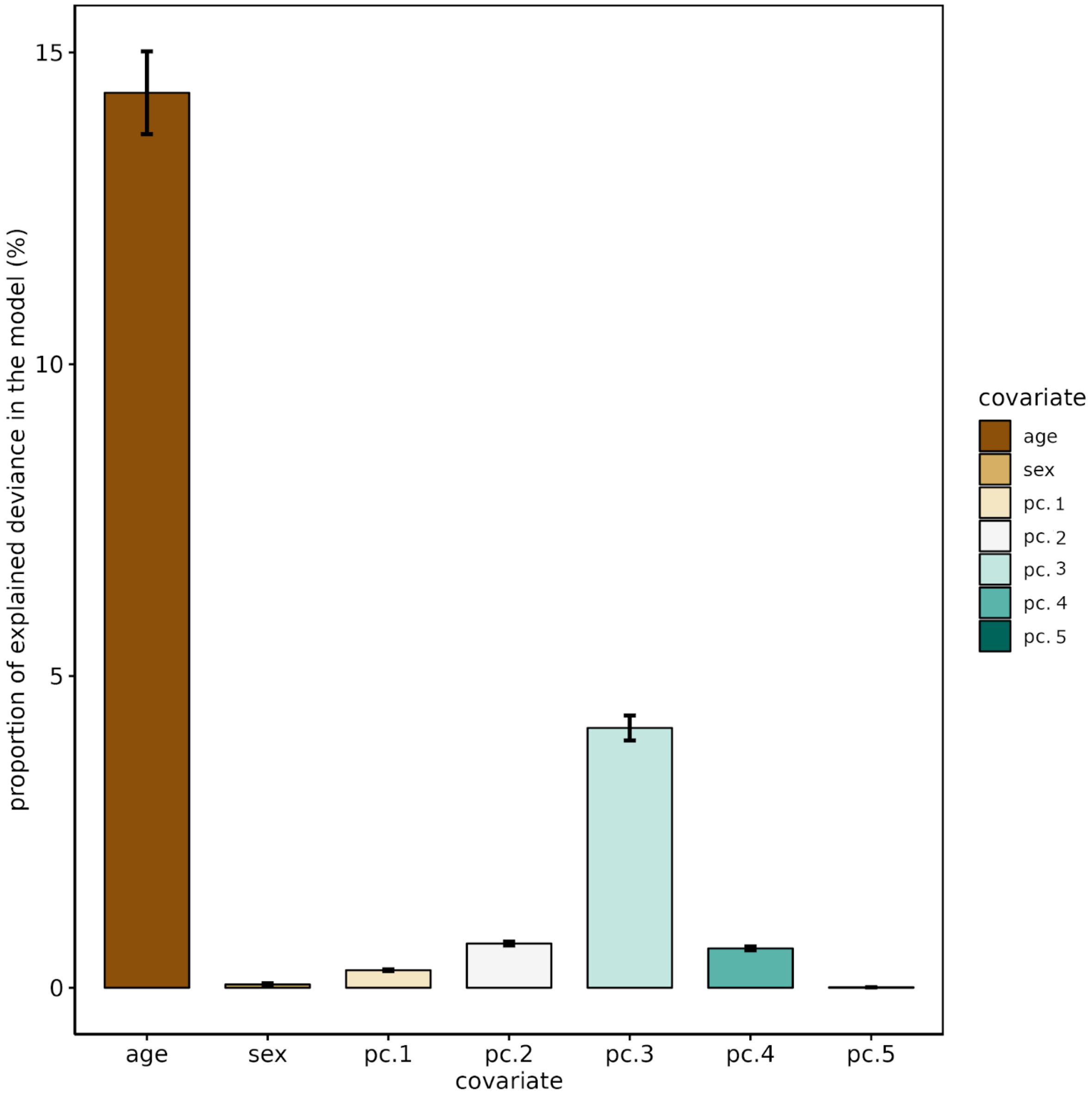

This bar plot displays the relative contributions of non-genetic covariates to the explained variance within the fully partitioned generalised linear model for height. The covariates included in the model are age, sex, and the first five principal components (PC1, PC2, PC3, PC4, and PC5). The y-axis represents the percentage of the model's explained variance, and the covariates are listed on the x-axis. Age is depicted as the predominant factor, explaining the highest percentage of variance, which is significantly more than any other single covariate. This suggests a strong age-related component in the phenotype's variation. The second most influential covariate is PC3, indicating a specific variance in the data captured by this principal component that is relevant to the phenotype. Sex, PC1, PC2, PC4, and PC5 show successively smaller contributions to the explained variance, implying that while they play a role in the phenotype's variation, their impact is less substantial compared to age and PC3. Error bars indicate the standard error for the proportion of variance explained by each covariate.

**Supplementary Figure 10:** Comparison of various non-genetic covariate interactions' contribution to height variance, as explained in the fully partitioned model.

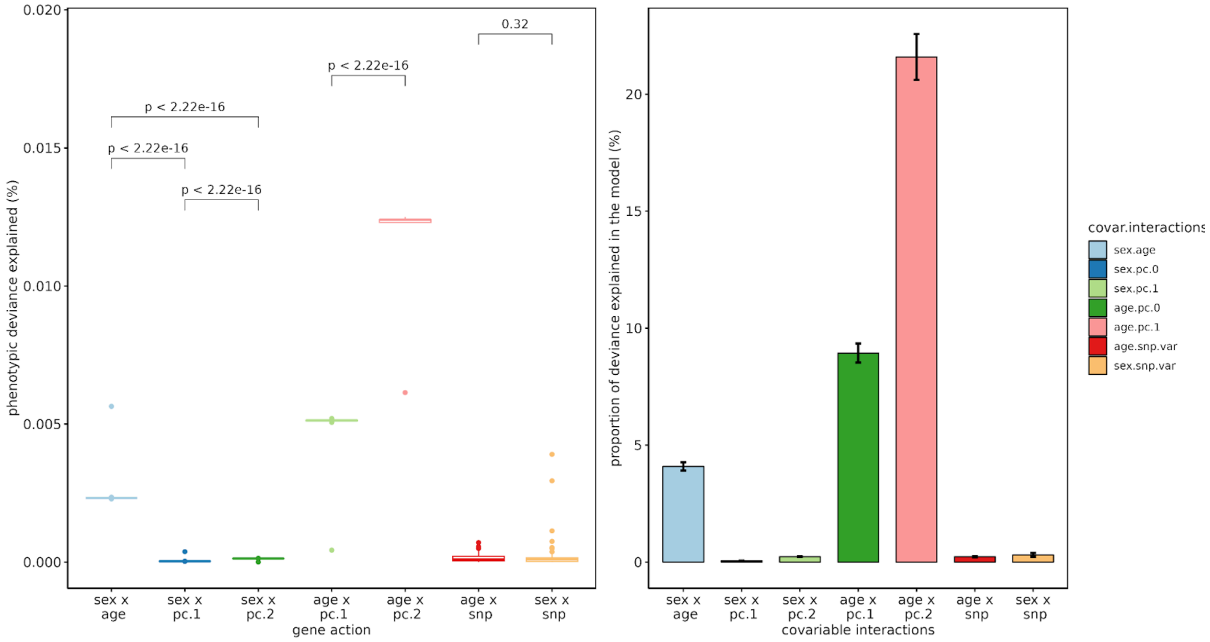

**Figure 10.a (left panel)- Total phenotypic variance explained by non-genetic covariate interactions:** The box plots show the total phenotypic variance explained by interactions between various non-genetic covariates: sex and age, sex and the first two principal components (PC1 and PC2), age and the first two principal components, and the interaction of sex and age with tested variants. **Figure 10.b (right panel) - Proportion of explained variance in the model by non-genetic covariate interactions:** The bar plot showing the proportion of the total variance explained by the model that is attributable to the same non-genetic covariate interactions. The bars are colour-coded according to the type of interaction, with the length of the bars reflecting the mean proportion of variance explained by each interaction term and error bars signifying the standard error. The sex by age and age by PC1 and PC2 interactions are shown to have a more pronounced impact on the phenotype. This underscores the different magnitudes of impact that various non-genetic covariate interactions have on height, with age-related interactions emerging as particularly influential.

**Supplementary Figure 11:** Trumpet plot illustrating the power for detecting additive-by-additive genetic interactions based on the analysis of 1,063 quasi-independent genome-wide significant (GWS) SNPs.

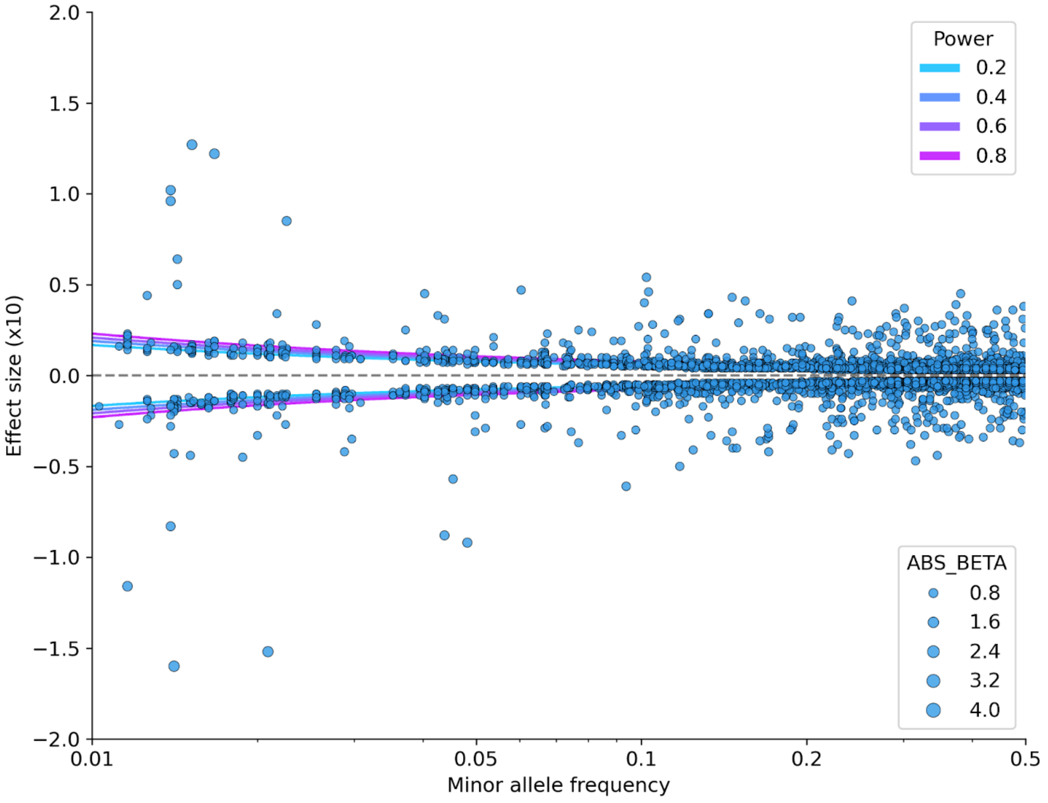

This trumpet plot visualises the results of a power analysis for detecting additive-by-additive genetic interactions affecting height, displayed across various minor allele frequencies (MAF) on the x-axis. The y-axis quantifies the strength of these genetic interactions, scaled by a factor of ten for clearer visualisation. Each dot represents a GxG interaction signal, with the size of the dot indicating the absolute effect size (ABS_BETA) of the interaction on height, expressed in standard deviations (s.d.) per minor allele.
